# Discovery of a pyrazolopyridine alkaloid that mitigates neuronal ER stress and age-related decline

**DOI:** 10.1101/2025.07.14.664722

**Authors:** Salinee Jantrapirom, Apiwat Sangphukieo, Natsinee U-on, Pattaporn Poonsawas, Wasinee Wongkumool, Ranchana Yeewa, Chansunee Panto, Puttachat Poound, Ester Zito, Alice Marrazza, Luca Lo Piccolo

## Abstract

Endoplasmic reticulum (ER) stress contributes to the pathogenesis of neurodegenerative and age-associated diseases, motivating the search for compounds that enhance ER-stress resilience. Modulation of ER-redox pathways, including those associated with the oxidase ERO1A, can attenuate maladaptive unfolded protein response (UPR) signaling and improve cellular stress tolerance. Here we developed an integrative discovery strategy to identify natural compounds that mitigate ER-stress-associated phenotypes across cellular and organismal models. Structure-informed virtual screening guided by ERO1A biology prioritized the pyrazolopyridine alkaloid S88. In human SH-SY5Y-derived neurons, S88 improved survival and reduced tunicamycin-induced ER-stress markers. In *Drosophila*, S88 ameliorated neuromuscular and locomotor phenotypes in a UBQLN2-associated ALS model and improved aging-related outcomes. Biochemical assays did not detect inhibition of ERO1A or radical scavenging activity by S88, indicating that its molecular target remains to be identified. Together, these findings identify S88 as a natural-product scaffold that enhances ER-stress resilience across neuronal and *in vivo* models.

**Graphical abstract:** 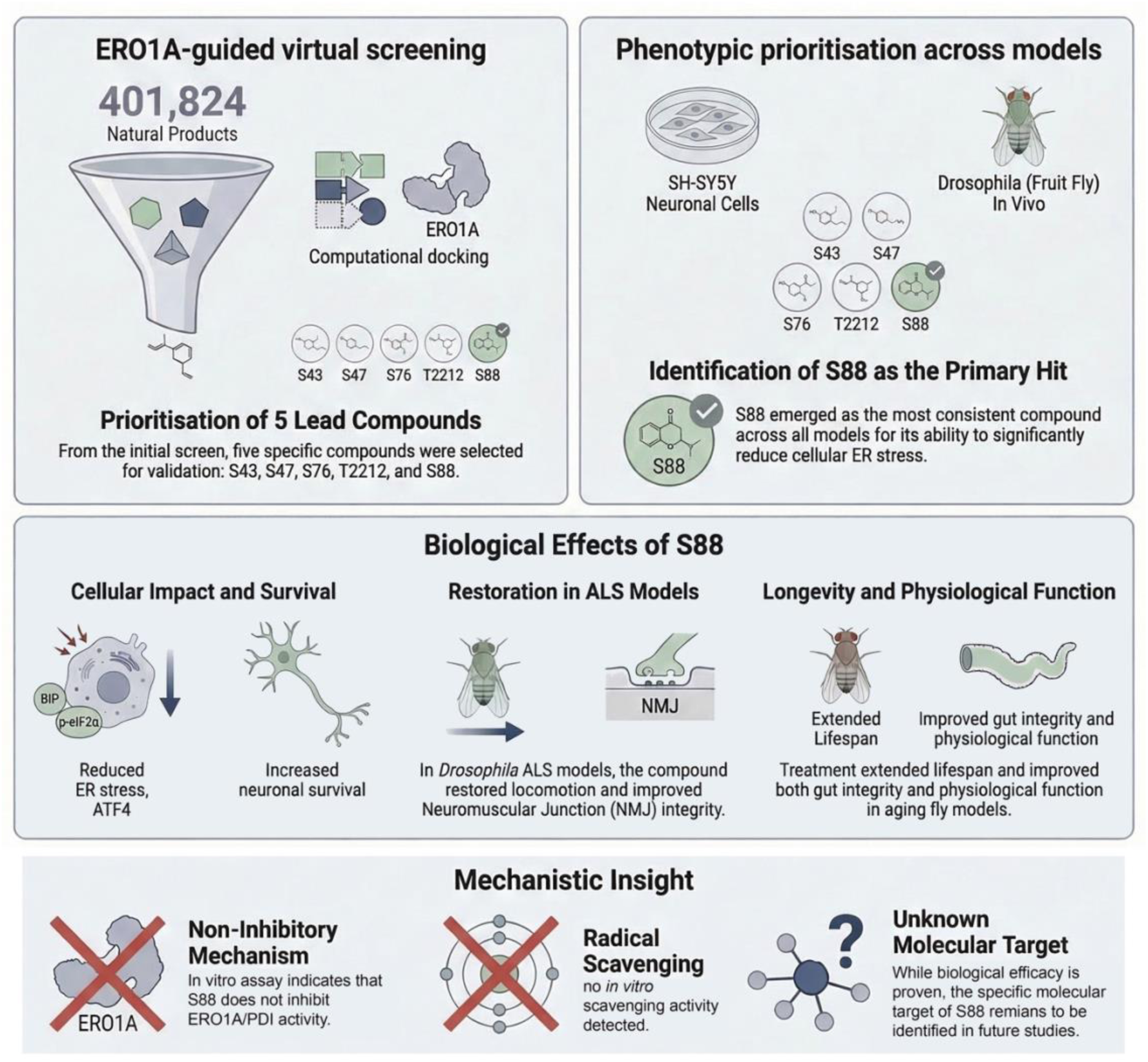

Structure-informed screening guided by ERO1A prioritized five natural products for functional validation across cellular and *Drosophila* models. The pyrazolopyridine alkaloid S88 consistently reduced ER-stress markers, improved neuronal survival, rescued locomotor and neuromuscular defects in an ALS model, and ameliorated aging phenotypes. The direct molecular target of S88 remains to be defined. Generated with assistance from the AI-based visualization tool NotebookLM and refined by the authors.

## 1. Introduction

Endoplasmic reticulum (ER) stress is a hallmark of several neurodegenerative diseases, including Amyotrophic Lateral Sclerosis (ALS), and plays a crucial role in aging-related neuronal dysfunction ^1-3^. The accumulation of misfolded proteins in the ER impairs cellular homeostasis, triggering the unfolded protein response (UPR). While the UPR initially functions as a protective mechanism to restore ER homeostasis, its prolonged activation may also contribute to neuronal cell death and disease progression. Thus, targeting ER stress pathways represents a promising therapeutic strategy for neurodegenerative diseases and age-related neuronal decline ^4^.

Among the regulators of ER stress, the Endoplasmic Reticulum Oxidoreductin 1 (ERO1) plays a crucial role in oxidative protein folding and maintaining ER redox poise. Its activity directly influences the burden of misfolded proteins in the ER, making it a critical factor in UPR activation ^5-7^. While ERO1 is essential in yeast and worms, partial inhibition enhances survival under ER stress and even extends lifespan in worms and *Drosophila* ^8-11^. In mammals, the presence of two isoforms, ERO1A and ERO1B, allows for functional flexibility, with loss-of-function mutations causing only mild defects ^12-14^. Collectively, these findings support the hypothesis that targeted inhibition of ERO1 could offer therapeutic benefits in conditions characterized by maladaptive UPR.

Notably, ERO1 inhibition has long been recognized as a potential anti-cancer strategy, given that many tumors exploit ER stress for survival and therapy resistance ^15-17^. More recently, we and others have demonstrated the therapeutic potential of targeting ERO1A in non-cancerous diseases. Suppressing ERO1A, either genetically or pharmacologically with EN460 or the pan-ER stress inhibitor TUDCA, has shown beneficial effects in a *Drosophila* model of UBQLN2-ALS as well as in both *in vitro* and *in vivo* models of SEPN1-related myopathy (SEPN1-RM) ^11,18^. These findings highlight ER redox regulation as a suitable target for therapeutic intervention in diseases characterized by chronic ER stress.

Despite this promise, the development of selective and well-tolerated small molecules that modulate ER redox pathways remains challenging. Existing ERO1A inhibitors such as EN460 exhibit limited selectivity and can interact with other flavin adenine dinucleotide (FAD)–dependent enzymes, including MAO-A, MAO-B, and LSD1 ^14,19^. FAD-dependent enzymes constitute a large family of flavoproteins that catalyze redox reactions using flavin adenine dinucleotide (FAD) as a cofactor. Because the architecture of FAD-binding domains is highly conserved across this enzyme class, compounds designed to target one flavoprotein may also interact with others, increasing the risk of off-target effects. ^20,21^. Previous virtual-screening efforts have yielded candidate ERO1A-directed chemotypes with improved predicted binding and *in vitro* activity. However, progress toward translation has been constrained by the scarcity of molecules that demonstrate robust, well-tolerated efficacy *in vivo*, and by the general difficulty of maintaining pathway-modulating activity under physiological conditions. These limitations highlight the need for discovery strategies that couple target-informed prioritization with early, cross-model functional validation to identify compounds that reproducibly ameliorate ER-stress-linked phenotypes in cells and organisms^22,23^.

Natural products represent a rich source of bioactive scaffolds for drug discovery due to their structural diversity and favorable drug-like properties ^24,25^. Here, we developed a multi-modal discovery pipeline combining structure-informed virtual screening with cross-platform functional assays to identify natural compounds capable of mitigating ER-stress-associated phenotypes. Using ERO1A-guided docking as a biologically informed prioritization framework, selected based on its central role in ER oxidative folding and stress-associated redox signaling, we screened 401,824 natural products from the COCONUT database and identified the heterocyclic pyrazolopyridine alkaloid CNP0410364 (hereafter S88) as a candidate molecule with robust biological activity.

S88 reduced UPR and ER-stress marker induction in multiple experimental systems and improved neuronal survival in human SH-SY5Y-derived neurons subjected to tunicamycin (Tm)-induced ER stress. In *Drosophil*a models of proteostasis dysfunction, including UBQLN2-associated ALS, S88 ameliorated neuromuscular and locomotor phenotypes and preserved functional performance during disease progression. Furthermore, S88 extended lifespan in a *Drosophila* model of accelerated aging induced by D-galactose. In addition, S88 improved lifespan, locomotor performance, and intestinal barrier integrity in naturally aged flies, indicating broader protective effects across both accelerated and physiological aging contexts.

Together, these findings identify S88 as a bioactive natural-product scaffold that mitigates ER-stress-associated phenotypes across neuronal and organismal models. Although ERO1A-guided docking served as the initial prioritization framework, the precise molecular target underlying S88 activity still remains to be defined. Nonetheless, these results highlight the therapeutic relevance of ER redox pathways linked to ERO1A and position S88 as a tractable small molecule for further target deconvolution and therapeutic development for ER-stress–associated neurodegenerative and age-related disorders.

## 2. Results

### 2.1 Phytochemical virtual screening prioritizes candidate ERO1A-targeting compounds with predicted selectivity over related FAD enzymes

A virtual screening of 401,824 phytochemical compounds was conducted to identify small molecules predicted to interact with ERO1A and potentially modulate ER stress pathways (Figure 1). We focused our primary docking screen on ERO1A because it is the stress-responsive mammalian isoform most commonly implicated in pathological ER-stress amplification, and it represents the closest human counterpart to the single *Drosophila* ortholog (Ero1L) that we previously identified as a genetic driver of ER-stress–associated phenotypes ^11^. Given the central role of ERO1A in oxidative protein folding and ER redox homeostasis, modulation of this pathway has been proposed as a strategy to mitigate maladaptive ER-stress signaling under pathological conditions.

**Figure 1.**
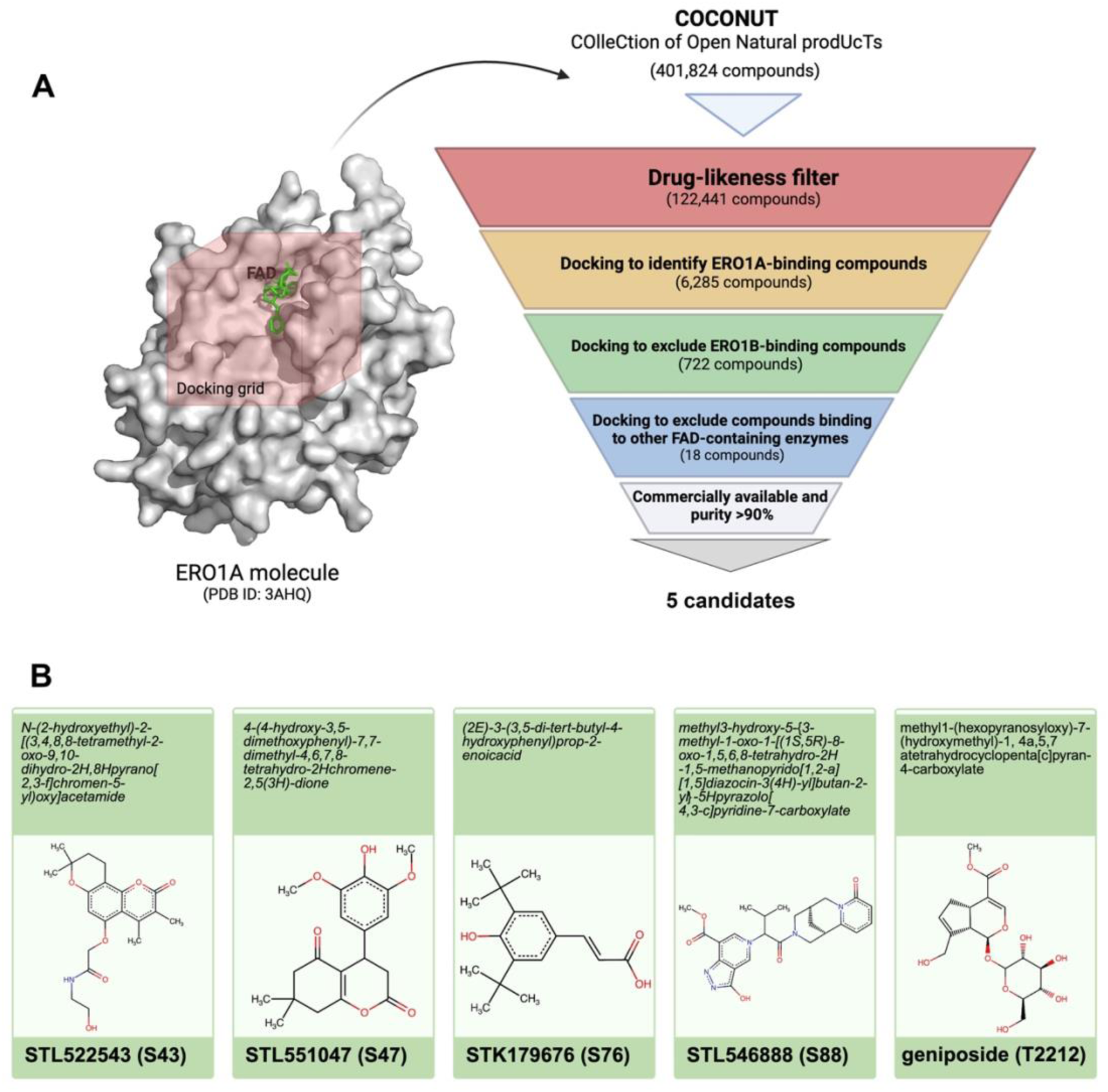
The workflow of the docking-based virtual screening process of phytochemical compounds targeting ERO1A. A) The 3D structure of human ERO1A (PDB ID 3AHQ) was prepared with a grid box defined around the binding pocket for docking simulations. Phytochemical compounds from the COCONUT database underwent virtual screening, starting with drug-likeness filtering. The remaining compounds were predicted in 3D and docked to ERO1A to assess binding affinities. Only compounds with a binding affinity higher than EN460, a known ERO1A inhibitor, were selected. Sequential docking against other FAD-containing enzymes (ERO1B, MAO-A, MAO-B, and LSD1) ensured specificity with only compounds showing minimal binding to these proteins being retained. Finally, only commercially available compounds with ≥90% purity were selected for *in vitro* validation and *Drosophila* model testing. B) Chemical structures of the five phytochemicals identified through screening and subsequently investigated *in vitro* and *in vivo* in this study.

Drug-likeness criteria based on the fraction of sp³ carbon atoms, lipophilicity, and molecular weight were applied as an initial filter to enrich the library for compounds with favorable physicochemical and pharmacokinetic properties, reducing the dataset to 122,441 candidates (30%). Because ERO1A is a FAD–dependent oxidoreductase that shares structural features with other FAD-containing enzymes, subsequent screening steps incorporated selectivity analyses to prioritize compounds predicted to preferentially interact with ERO1A over related FAD enzymes.

Next, these compounds were screened for their binding affinity to ERO1A, retaining only those with stronger affinity than EN460 (-8.6 kcal/mol), a known ERO1A inhibitor. This step narrowed the selection to 6,825 compounds (6%).

To further assess predicted selectivity, we compared the docking affinities of these compounds against other FAD-containing enzymes, reflecting the highly conserved nature of FAD-binding sites. Binding affinity to ERO1B, the paralog of ERO1A, was first evaluated; 722 compounds (∼11%) exhibited stronger predicted affinity for ERO1A than for ERO1B. These compounds were then further screened against additional FAD-containing enzymes, including MAO-A, MAO-B, and LSD1. The binding energies of known inhibitors, harmine (−8.7 kcal/mol) for MAO-A, safinamide (-9.3 kcal/mol) for MAO-B, and CC-90011 (-9.9 kcal/mol) for LSD1, were used as reference thresholds for favorable binding. Most compounds exhibited equal or stronger predicted binding to these enzymes than their respective inhibitors, leaving 18 compounds (3%) predicted to preferentially interact with ERO1A under our docking conditions (Supplementary Tables 1-3). Of these 18 compounds, only 8 were available for synthesis, with 5 commercially purchasable at ≥90% purity (Supplementary Table 4). Compounds CNP0392367 and CNP0168092 (hereafter referred to as S43 and S47, respectively) and geniposide were purchasable with a perfect match, while CNP0079545 and CNP0410364 (hereafter referred to as S76 and S88, respectively) were available only as isomers (Supplementary Table 4).

### 2.2 Selected phytochemicals show minimal cytotoxicity and improve survival in an ERO1L-sensitized *Drosophila* model

Before evaluating their functional activity, we first assessed the potential cytotoxicity of the selected phytochemicals in human cells in order to exclude compounds with intrinsic toxicity that could confound downstream phenotypic analyses. Primary human dermal fibroblasts (HDF), HepG2, and SH-SY5Y cells were exposed to increasing concentrations of each compound for 24 hours (Supplementary Figure S1) or 48 hours (Figure 2).

**Figure 2.**
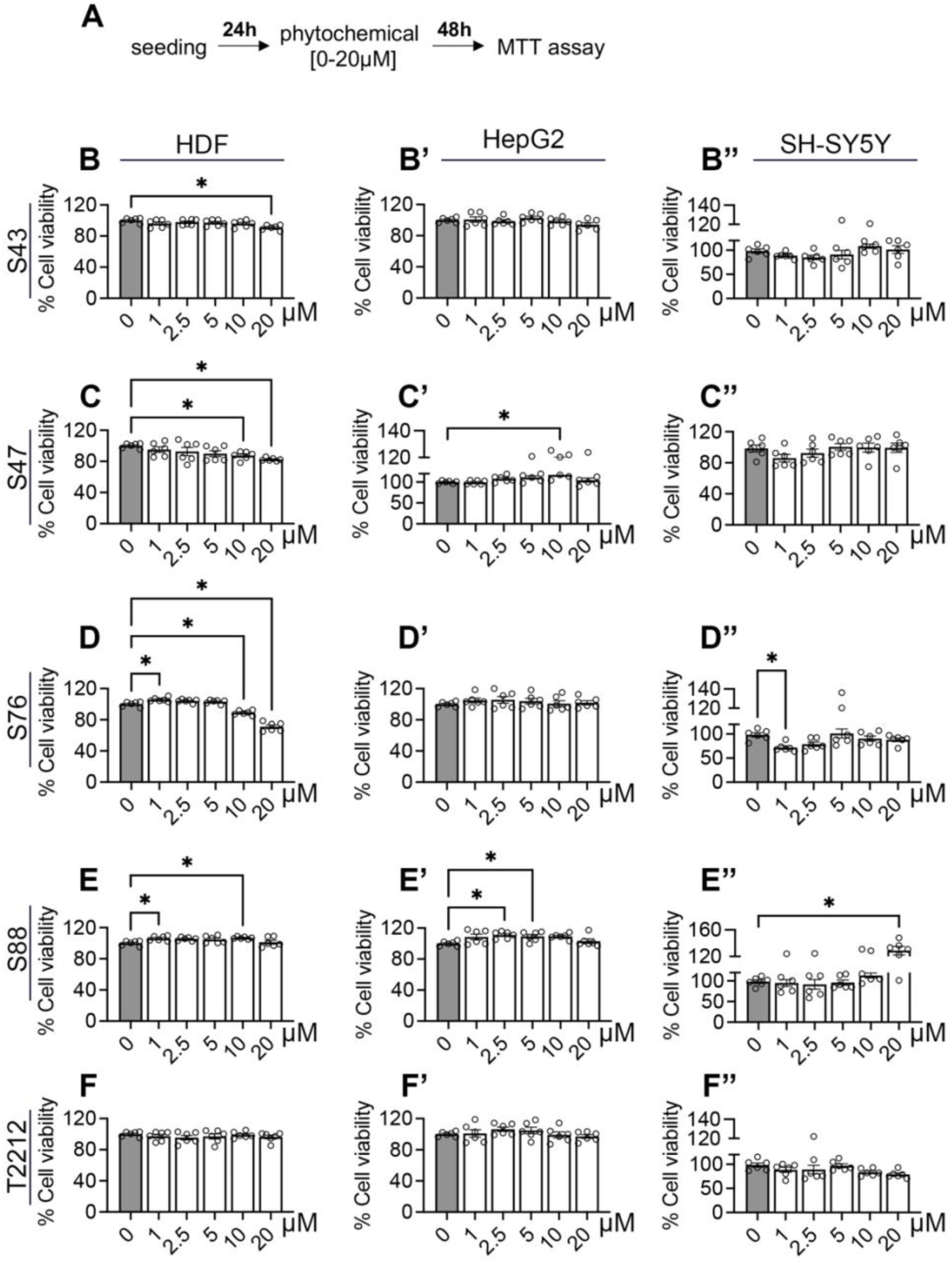
The administration of S88 or T2212 does not induce toxicity in primary cells or cell lines. A) Experimental design scheme. B-F’’) Cell viability of human-derived fibroblast (HDF) (B-F), Hep2G (B’-F’), and SH-SY5Y (B’’-F’’) cells after 48-hour (h) incubation with increasing concentrations of identified phytochemicals (0 to 20 µM). Cell viability was assessed by MTT assay and normalized to vehicle-treated controls (0.1% DMSO). Bars represent mean ± SD, and individual values correspond to independent replicates (n = 6). The data illustrate the effects of progressive concentration increases on cell viability across the different cell types. Statistical significance was assessed by one-way ANOVA followed by Dunnett’s T3 multiple-comparisons test, with comparisons made to the vehicle control (*p < 0.05). The data related to cell viability after 24h phytochemical treatment are shown in Supplementary Figure S2.

S43, S47, and S76 showed detectable cytotoxicity in primary fibroblasts at the highest doses after 24 h (Supplementary Figure S1B, C), with S47 exhibiting a time-dependent increase in toxicity in HDF cells (Figure 2C). In contrast, neither S88 nor T2212 showed measurable toxicity in HDF cells at any incubation time tested (Figure 2E–F, Supplementary Figure S1D–E). Across all compounds, cytotoxic effects were limited and primarily observed in primary fibroblasts at higher concentrations, whereas HepG2 and SH-SY5Y cells displayed minimal sensitivity under the conditions tested. These results indicate that most compounds are well tolerated in human cells within the concentration ranges used in subsequent assays. We next evaluated whether the selected phytochemicals could improve organismal outcomes in an ER-stress-sensitized *in vivo* context. To this end, we utilized the shortened lifespan of *nSyb>ERO1L* flies, which results from neuronal overexpression of ERO1L, the *Drosophila* ortholog of mammalian ERO1A ^11^. Survival analysis was performed on these flies, both with and without phytochemical treatment, to determine whether the compounds could mitigate the lifespan defect associated with elevated ERO1L activity. All five phytochemicals significantly improved survival relative to vehicle-treated *nSyb>ERO1L* flies, as reflected by increased median lifespan (Figure 3 A-E).

**Figure 3.**
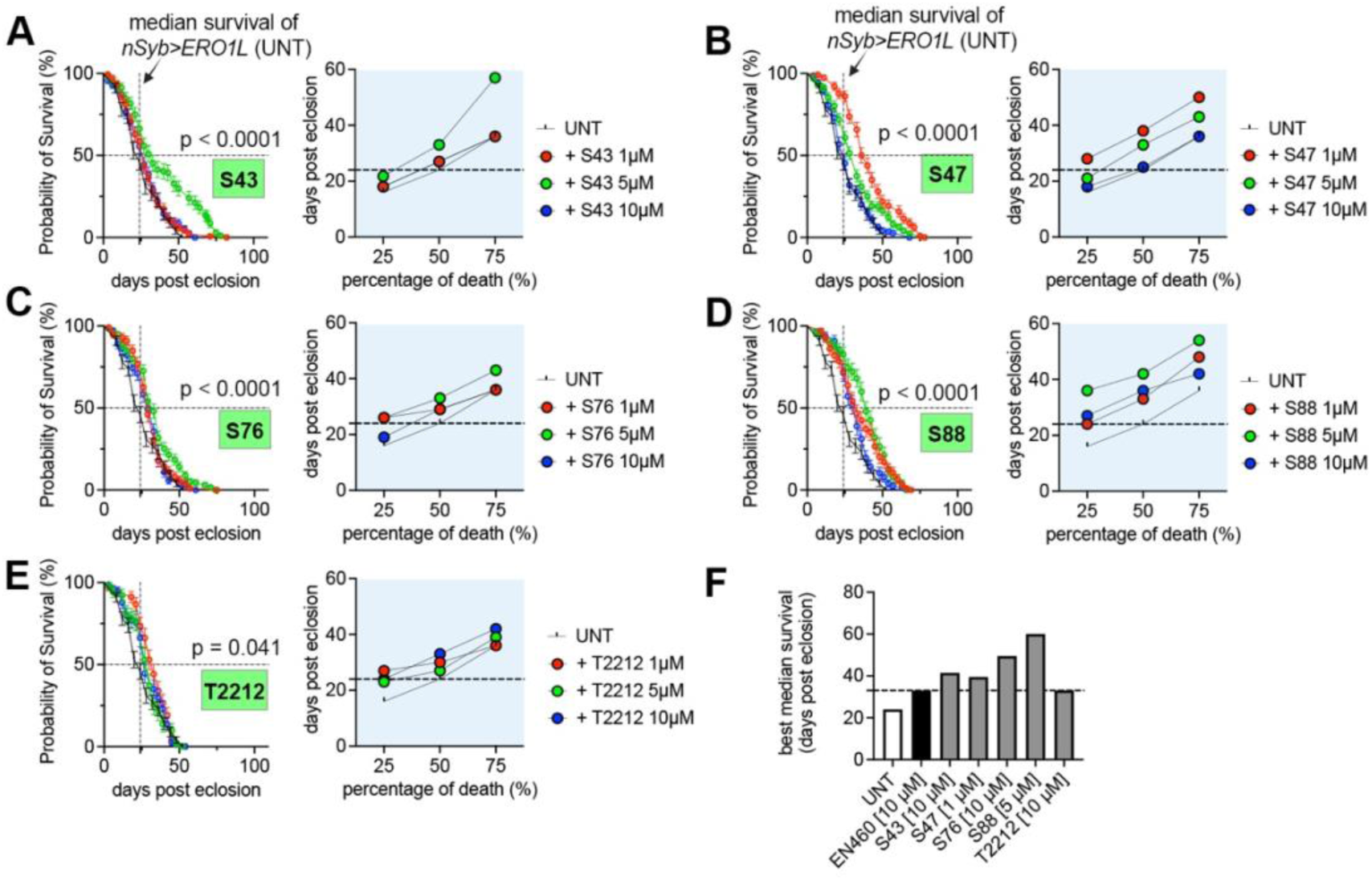
Identified phytochemicals attenuate ERO1-associated stress phenotypes *in vivo*. (A–E) Survival curves of pan-neuronal (*nSyb-GAL4*) ERO1L-overexpressing flies treated with increasing concentrations (1, 5, or 10 µM) of S43 (A), S47 (B), S76 (C), S88 (D), and T2212 (E), or maintained on standard food containing vehicle only (untreated control, UNT; 0,025% DMSO). Lifespan experiments were performed in parallel under identical environmental conditions, with a shared untreated control cohort included in the same experimental batch. For clarity of visualization, survival curves are presented in separate panels grouped by phytochemical treatment, while the untreated control curve is shown in each panel as the common reference population. Each phytochemical condition included n = 100 flies, and the shared untreated cohort included n = 389 flies. Statistical significance was determined using a log-rank (Mantel–Cox) test applied to the full survival dataset, comparing untreated flies (UNT) with those treated with 1, 5, or 10 µM compound. Insets show the quartile percentage of mortality at each tested dose. The median survival of untreated *nSyb>ERO1L* flies was 24 days; a dotted line is overlaid to facilitate comparison across treatments. (F) Bar plot showing the highest median survival observed for *nSyb>ERO1L* flies following phytochemical treatment. The *in vivo* activity of the identified phytochemicals is compared with EN460, whose full survival curve is shown in Supplementary Fig. S1.

Although the magnitude of the effect varied depending on compound and dose, these data demonstrate that the identified molecules exert *in vivo* activity in an ERO1L-sensitized context, recapitulating a protective effect similar to EN460 under these conditions (Figure 3 F, Supplementary Figure S2).

S43 and S88 were the most effective in extending lifespan, with 75% of the flies surviving at 60 and 50 days post-eclosion at 5 µM, respectively (Figure 3 A and D). Notably, a 5 µM dose of S43 showed the most pronounced benefit, as the survival probability slope dramatically decreased compared to untreated or other doses, suggesting a potential time-dependent pro-longevity effect that may complement its ERO1L inhibitory activity (Figure 3 A). On the other hand, S47 showed the greatest benefit at the lowest concentration (1 µM), implying potential toxicity related to its metabolism at higher concentrations (Figure 3 B).

Both S76 and T2212 extended lifespan to a lesser degree (Figure 3C and E). S76 demonstrated similar effects at 1, 5, or 10 µM, with a slightly higher impact at 5 µM (Figure 3C). T2212 was most effective at 10 µM (Figure 2E).

Finally, S88 extended lifespan across all tested doses, with the most pronounced benefits observed during early adulthood. Consistent with this, the time to 25% mortality was increased in S88-treated flies relative to vehicle-treated controls and the other phytochemical conditions. S88 exhibited particularly strong effects at both 1 µM and 5 µM (Figure 3D).

Together, these results demonstrate that several prioritized phytochemicals exhibit *in vivo* activity in an ER-stress-sensitized model while displaying limited cytotoxicity in human cells.

### 2.3 S88 and T2212 alleviate ER stress markers and enhance survival in Tm-stressed neuronal cells

To determine whether the prioritized phytochemicals could modulate ER-stress responses, we examined their effects on ER-stress marker induction and cell survival under tunicamycin (Tm) challenge. To this end, we initially focused on SH-SY5Y neuroblastoma cells, a widely used neuronal model for studying ER-stress-associated mechanisms relevant to neurodegenerative processes.

We first established an optimized model for ER stress induction by treating undifferentiated SH-SY5Y cells with 1 µM tunicamycin (Tm) and monitored cell viability along with the expression of key ER stress markers over time (Supplementary Figure S3A). As expected, Tm exhibited a time-dependent cytotoxic effect, with a significant reduction in cell viability after 48h (Supplementary Figure S3 B-D). Interestingly, ER stress markers peaked at 24h before declining, suggesting an early transcriptional response followed by progressive cellular damage (Supplementary Figure S3 E). Based on these findings, we chose a 48h Tm incubation period for subsequent experiments.

To assess the beneficial effects of phytochemicals, we employed a stepwise treatment approach, administering the compounds after the onset of cellular stress. This involved an initial 24h exposure to Tm, followed by a 24h treatment with each phytochemical at either 10 or 20 µM (Figure 4A and Supplementary Figure S4A). Cell viability and ER stress markers were evaluated at the end of the treatment period.

**Figure 4.**
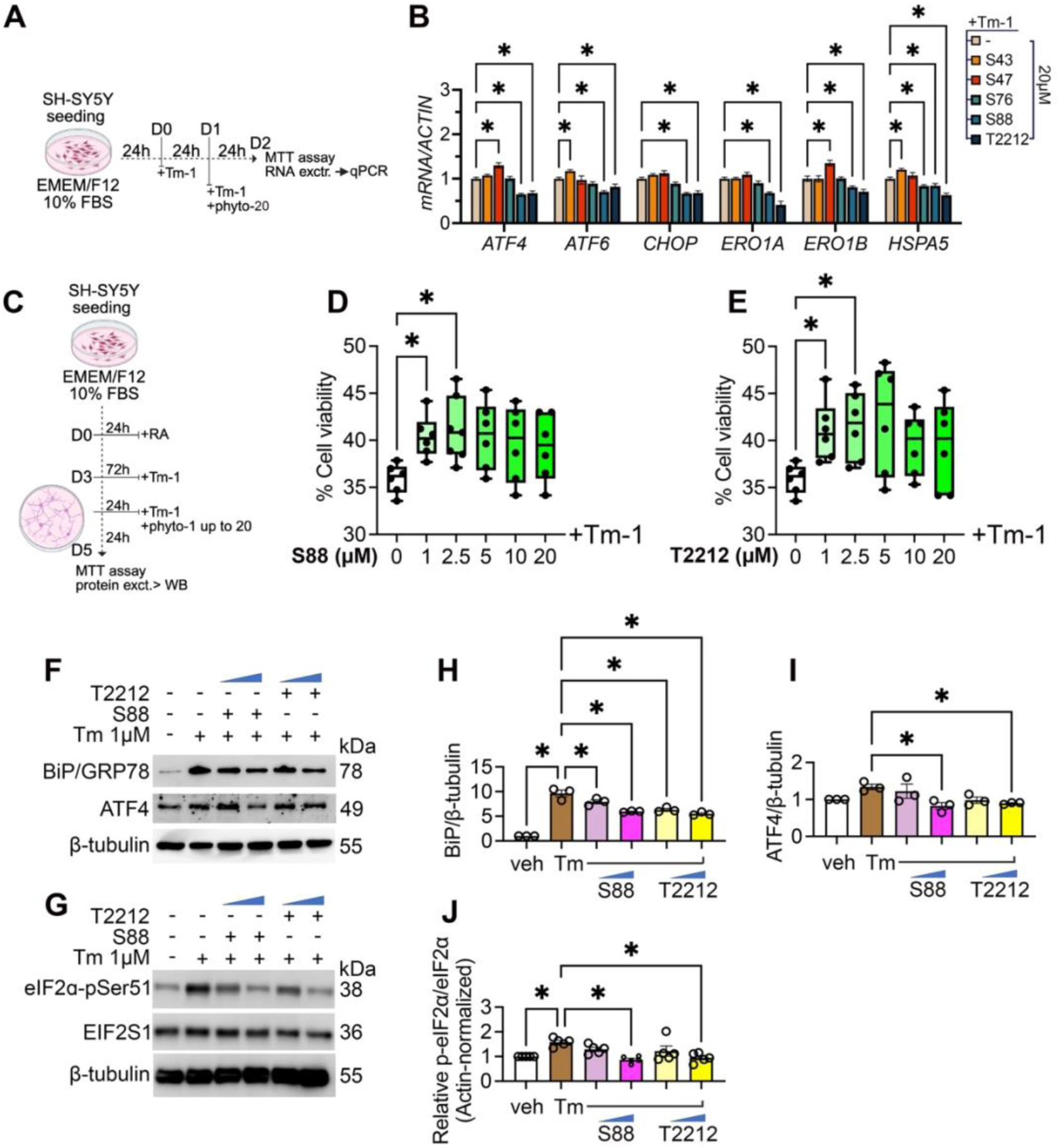
S88 and T2212 enhance viability and reduce ER stress markers in Tm-stressed SH-SY5Y cells. A) Experimental design with undifferentiated SH-SY5Y cells. B) Expression levels of ER stress markers in Tm 1 µM (Tm-1)-stressed SH-SY5Y cells after 24 hours (h) phytochemical (phyto) administration at 20 µM. Dose- and time-dependent response of Tm-1-stressed undifferentiated SH-SY5Y cells is shown in Supplementary Figure S3. C) Experimental design with Retinoic acid (RA)-differentiated SH-SY5Y cells. D-E) Cell viability of SH-SY5Y cells Tm-1 stressed for 24h, followed by 24h treatment with increasing concentration of either S88 (D) or T2212 (E) in the continued presence of Tm-1. Cell viability was measured using the MTT assay and compared to vehicle (0.1% DMSO) controls. (F) Representative immunoblots showing BiP/GRP78 and ATF4 and (G) p-eIF2α and eIF2α (EIF2S1). (H–I) Quantification of BiP/GRP78 levels normalized to β-tubulin and expressed relative to control. (J) Relative p-eIF2α/eIF2α levels (Actin-normalized). Data are shown as individual values with mean ± SD. Statistical significance was determined by one-way ANOVA followed by Sidak’s multiple comparison test. *p < 0.05.

Among the five tested compounds, only S88 and T2212 consistently improved cell viability under these experimental conditions while significantly reducing all measured ER stress markers (Supplementary Figure S4B, C and Figure 4B). Although S76 produced the largest increase in cell viability (Supplementary Figure S4B), this effect was not accompanied by a general reduction in ER stress markers. Only *HSPA5* levels were lower compared to the untreated Tm-stressed control when S76 was provided at 20 µM (Figure 4B). Similarly, S43 improved cell survival but at the same time led to an increase in most ER stress markers, except for *ATF4*, which remained unchanged (Supplementary Figure S4C and Figure 4B).

Taken together, these results identified S88 and T2212 as the only compounds that consistently coupled improved neuronal survival with reduced ER-stress/UPR marker induction across assays. Although several additional compounds increased viability under tunicamycin challenge, these effects were not accompanied by a corresponding reduction in ER-stress markers, making it unclear whether the observed protection reflected ER-stress attenuation or downstream compensatory responses. Because suppression of ER-stress signaling under neurotoxic conditions is associated with neuronal protection, S88 and T2212 were selected for further characterization.

Next, we validated our findings in mature neurons and sought to determine an appropriate Tm dose that induces ER stress without causing excessive cell death. SH-SY5Y cells were differentiated with RA (Supplementary Figure S5A), and as expected, neurite elongation was accompanied by a progressive increase in key neuronal maturation markers (Supplementary Figure S5B, C). Meanwhile, Tm exhibited a time- and dose-dependent cytotoxic effect, with significant cell death observed after 48h of exposure to 10 µM Tm (Supplementary Figure S6). Based on these findings, we implemented a 24h preconditioning step with 1 µM Tm (Tm-1) in 3-day RA-differentiated SH-SY5Y cells, followed by a 24h intervention with S88 or T2212 in the continued presence of Tm-1 (Figure 4C). Here, both phytochemicals improved neuronal survival under ER stress in a dose-dependent manner, with maximal protection observed at 2.5 µM and a decline in efficacy at higher concentrations (Figure 4D and E).

We next examined canonical markers of ER stress under these experimental conditions and found that treatment with 2.5 µM S88 or T2212 significantly reduced the levels of PERK, BiP, ATF4, and phosphorylated eIF2α (eIF2α-pSer51) compared to Tm-stressed neurons without phytochemical treatment (Figure 45F-J).

Taken together, these results indicate that both S88 and T2212 enhance neuronal survival under Tm-induced ER stress, likely through partial modulation of the UPR signaling pathway. These effects may reflect ERO1A modulation and/or additional targets within ER proteostasis pathways. These findings further support their candidacy as neuroprotective agents and prompted us to further investigate the mechanisms of action of S88 and T2212 in a more physiologically relevant context.

### 2.4 S88 and T2212 improve early neuronal health in UBQLN2^ALS^ models, but only S88 preserves locomotor function throughout disease progression

Next, we turned to a *Drosophila* model of neurodegeneration, which offers a complex *in vivo* system recapitulating key features of ER stress-associated neuronal pathology ^26-28^. We previously demonstrated that pharmacological inhibition of ERO1L with EN460 can improve pathophysiology of flies expressing the familial ALS-associated UBQLN2^P497H^ variant (UBQLN2^ALS^)^11,29^, a model in which chronic ER stress and maladaptive UPR activation are key pathogenic features ^30-32^. Notably, this rescue effect is specifically linked to ERO1L suppression, as we further showed that pan-neuronal *ERO1L* knockdown also restores locomotor function in this UBQLN2^ALS^ model (Supplementary Figure S7).

To assess the effects of S88 and T2212, increasing concentrations were administered to the parental cross and maintained throughout the development of the F1 generation (Figure 5A). Locomotor function was evaluated in larvae expressing UBQLN2^ALS^ in all neurons and compared to untreated controls (vehicle, veh). To establish a genetically matched baseline for normalization, we included an *elav>lacZ* control in which the same neuronal driver expresses an inert reporter rather than UBQLN2^ALS^. This design controls for driver background and transgene expression context. Moreover, elav>lacZ is compatible with NMJ morphology analyses because it does not introduce fluorescent reporters that could interfere with the HRP-FITC and Dlg immunostainings used to quantify synaptic architecture. A dose-dependent improvement in locomotion was observed (Figure 5B-C), alongside significant NMJ health benefits, including increased branch length and a higher number of synaptic boutons (Figure 5D-F, Supplementary Figure S8 and S9). Notably, S88 was effective at both 1 µM and 5 µM, whereas T2212 showed benefits only at the higher dose of 10 µM (Figure 5B-F). At these effective doses, larval locomotion and NMJ parameters were not statistically different from the CTRL (*elav>LacZ*) reference (Figure 5B-F), indicating functional restoration to baseline levels under the conditions tested.

**Figure 5.**
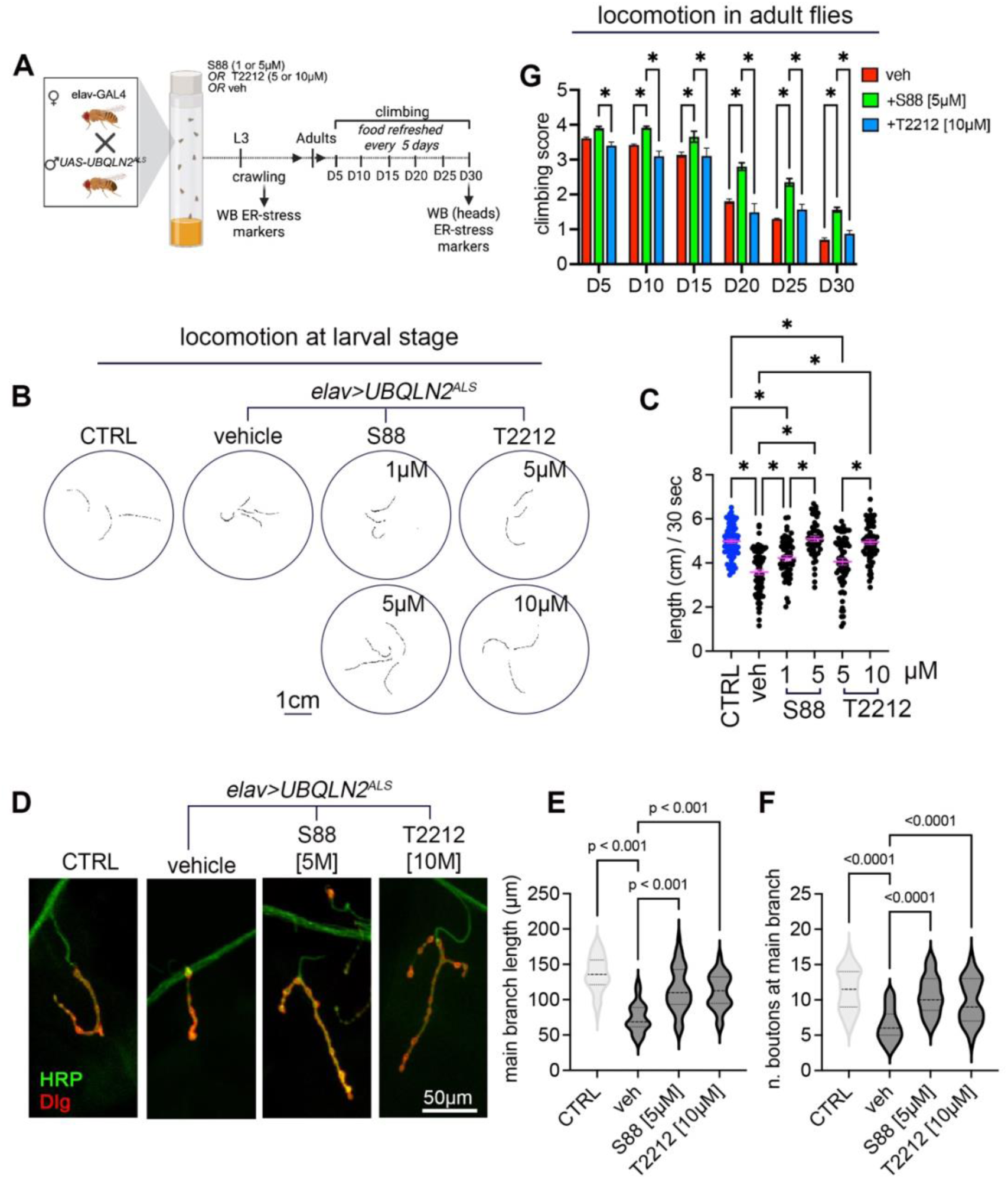
S88 and T2212 improve early neuronal health in UBQLN2^ALS^ models, but only S88 sustains late-stage locomotor benefits. A) Experimental design. B) Larval movement was quantified by measuring the total distance travelled within 30 seconds (sec), presented as a scatter plot with geometric mean. A genetically matched control line (CTRL; *elav>LacZ*) is included for baseline comparison. (C) Representative locomotor trajectories for each genotype tested. D) Type I motor neurons (Ib MNs4) innervating muscle 4 in larvae expressing UBQLN2^ALS^ under elav-GAL4 (*elav> UBQLN2^ALS^*) were immunoassayed for the presynaptic membrane marker horseradish peroxidase (HRP, green) and the postsynaptic marker Discs large (Dlg). CTRL (*elav>LacZ*), vehicle-treated (veh), and compound-treated larvae are shown. Scale bars: 50 μm. Single fluorescent channels and additional representative NMJ are shown in Supplementary Figure S8 and S9. (E–F) Quantification of NMJ morphology, including (E) branch length and (F) synaptic bouton number, measured using ImageJ (v1.53k). Data are presented as violin plots showing the distribution of individual NMJs. (G) Locomotor functions in adult *elav> UBQLN2^ALS^* flies were assessed using a climbing assay. Flies were either untreated (veh, n = 127) or treated with varying concentrations of S88 (n = 103) or T2212 (n = 112). Climbing performance was recorded at 5, 10, 15, 20, 25, and 30 days post-eclosion, with scores compared across age-matched groups. Statistical analysis was conducted using one-way ANOVA followed by Dunnett’s test for crawling and NMJ assessments, while two-way ANOVA with Tukey’s multiple comparisons test was applied for the climbing assay. Statistical significance was considered at *\*p* < 0.05.

To determine whether these improvements persisted into adulthood, we assessed locomotor performance in adult flies at different time points during aging (Figure 5G, Supplementary Figure S10).

In the climbing assay, S88 significantly improved motor performance compared to vehicle-treated UBQLN2^ALS^ flies. Notably, between D5 and D15, S88-treated flies were not significantly different from age-matched healthy reference controls (*elav>EGFP* and *elav>DsRed*), indicating normalization of locomotor function during early adulthood. At later ages (>D15), climbing performance remained improved relative to untreated mutants but declined compared to healthy controls, consistent with age-dependent attenuation of efficacy (Fig. 5G; Supplementary Figure S10). In contrast, T2212 did not sustain adult motor improvement, as performance progressively declined in parallel with vehicle-treated UBQLN2^ALS^ flies (Fig. 5G). Collectively, these results show that S88, but not T2212, improves adult locomotor function, with age-dependent attenuation of efficacy.

To determine whether the locomotor improvements observed in UBQLN2^ALS^ *Drosophila* models following phytochemical treatment were associated with ER stress modulation, we analyzed UPR markers and stress-responsive gene expression across disease stages and tissues (Figure 6A, Supplementary Figure S11A). Immunoblot analysis revealed that both S88 and T2212 reduced phosphorylated eIF2α (eIF2α-pSer51) at larval and adult stages (D30), indicative of attenuated ER stress (Figure 6B, D, F, and H). However, ATF4 (Crc in *Drosophila*) protein levels remained unchanged in S88-treated flies, despite reduced p-eIF2α (Figure 6C, G), suggesting an alternative mechanism sustaining ATF4 expression. Given reports of p-eIF2α–independent ATF4 regulation via mTORC1 and 4E-BP1 ^33,34^, we assessed mTORC1 activation by probing p70S6K and 4E-BP1 phosphorylation. Neither showed changes upon S88 treatment at any time point (Supplementary Figure S11), arguing against mTORC1 involvement.

**Figure 6.**
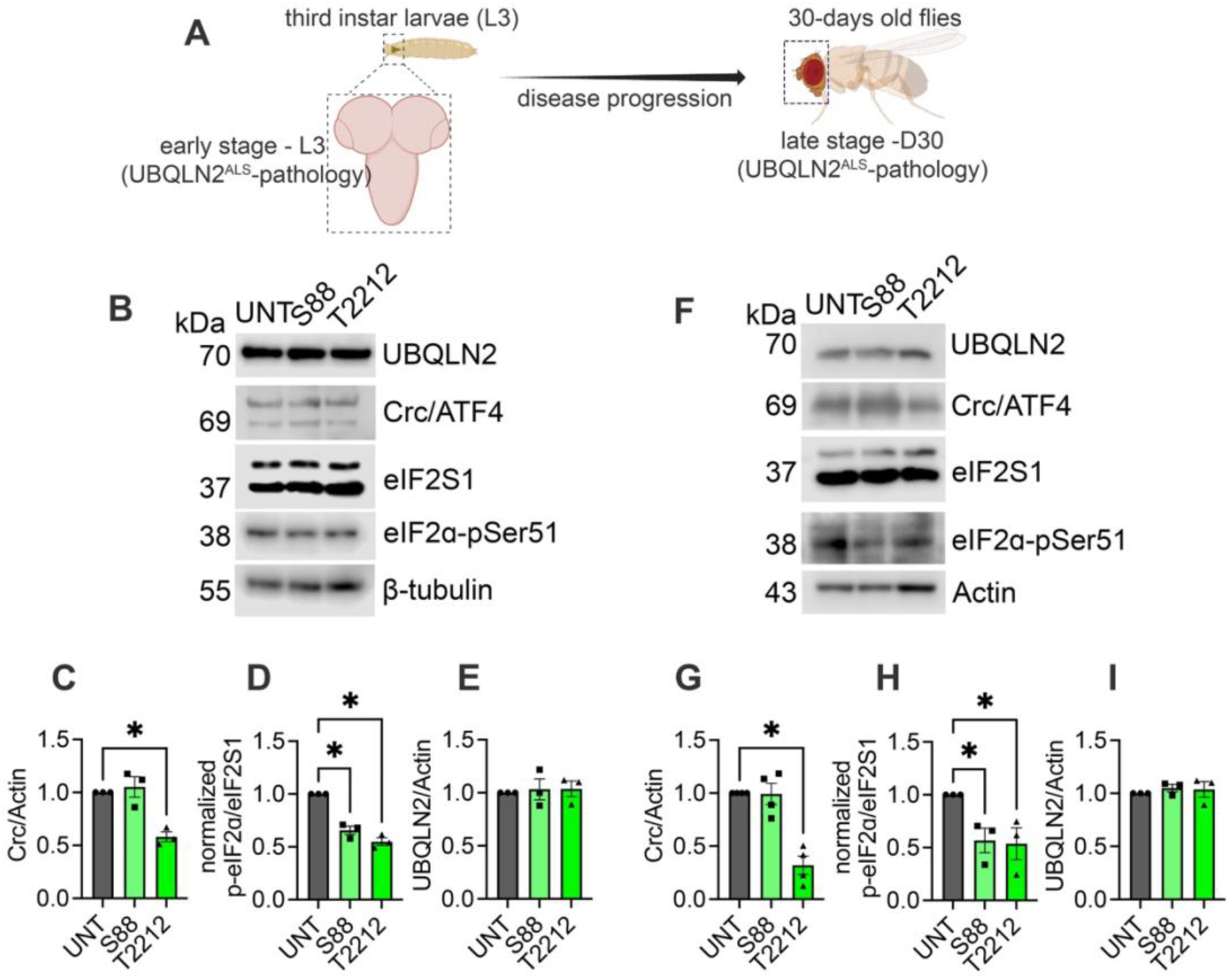
Efficacy of S88 in late-stage UBQLN2^ALS^ flies is associated with reduced p-eIF2α but sustained ATF4 levels. A) Schematic representation of disease progression and time points for ER stress marker analysis. (B, F) Representative immunoblots from larval brains (early disease stage; B) and heads of 30-day-old (D30) flies (late disease stage; F) expressing UBQLN2ALS and treated with S88 or T2212 or left untreated (UNT). (C–E, G–I) Quantification of the indicated proteins from early-stage larval brains (C–E) and late-stage adult heads (G–I). ER stress marker levels were compared with age-matched untreated controls. UBQLN2 levels were also assessed across treatment groups. Actin or β-tubulin served as loading controls. Each point represents an independent biological replicate; bars indicate mean ± SD. Statistical significance was assessed using one-way ANOVA followed by Šidák’s multiple-comparisons test relative to untreated controls (p < 0.05).

Transcriptional profiling uncovered distinct tissue- and stage-specific responses. In fly heads, both phytochemicals reduced multiple ER stress and cytoprotective gene transcripts at day 15 (D15-H), but this response waned by day 30 (D30-H), except for sustained *Sip3* downregulation in S88-treated flies (Supplementary Figure S12B, D). In decapitated fly bodies (carcass; non-cephalic tissues), transcriptional responses were more heterogeneous, consistent with a composite signal arising from multiple peripheral tissues following oral compound administration. *ATF4* mRNA was reduced at D15 but trended upward by D30, whereas *ATF6* levels were consistently decreased across compounds and time points. Additional stress-associated markers (*Pdi*, *Erp60*, *Hsp22*, and *Sip3*) exhibited dynamic, compound-specific regulation, with S88 producing the most consistent suppression of *Sip3* and *Hsp22* (Supplementary Figure S12C, E). Because UBQLN^ALS^ is expressed in neurons, carcass qRT–PCR is not expected to directly reflect the primary neuronal ER-stress response. Rather, these analyses assess whether stress-associated transcription becomes detectable in peripheral tissues during disease progression and whether orally administered compounds modulate these systemic responses, potentially contributing to organismal benefit.

Together, these data suggest that both S88 and T2212 mitigate ER stress during early disease. However, only S88 induces a residual stress-adaptive transcriptional program that persists into late-stage neurodegeneration, coupled with sustained ATF4 protein levels via a mechanism independent of both eIF2α phosphorylation and mTORC1.

### 2.5 S88 improves survival and age-associated functional decline in *Drosophila* aging models

Aging is marked by a progressive loss of proteostasis and heightened vulnerability to ER stress ^4,35,36^, processes that intersect with ER redox homeostasis and oxidative protein folding networks in which ERO1A has been implicated ^37,38^. Building on the neuroprotective effects of S88 and T2212 under both environmental (Tm-induced) and genetic (UBQLN2^ALS^-driven) ER stress and motivated by the broader link between ER-stress modulation and organismal resilience, we next evaluated whether these compounds could mitigate aging-associated functional decline *in vivo*. We first employed a *Drosophila* model of accelerated aging induced by chronic D-galactose (D-GAL) exposure, which recapitulates key features of aging, including oxidative stress, mitochondrial dysfunction, and reduced lifespan^39,40^. Following dose-dependent evaluation (Supplementary Fig. S13), 25 mM D-GAL was chosen for subsequent experiments as it induces a robust but not overly severe aging phenotype, allowing sensitivity to detect protective effects. Under these conditions, S88 significantly extended lifespan at both 1 µM and 5 µM (S88-1 and S88-5) compared to untreated controls (Fig. 7A). In contrast, T2212 did not confer a survival benefit at either tested dose, indicating limited efficacy in this accelerated aging context (Fig. 7B).

**Figure 7.**
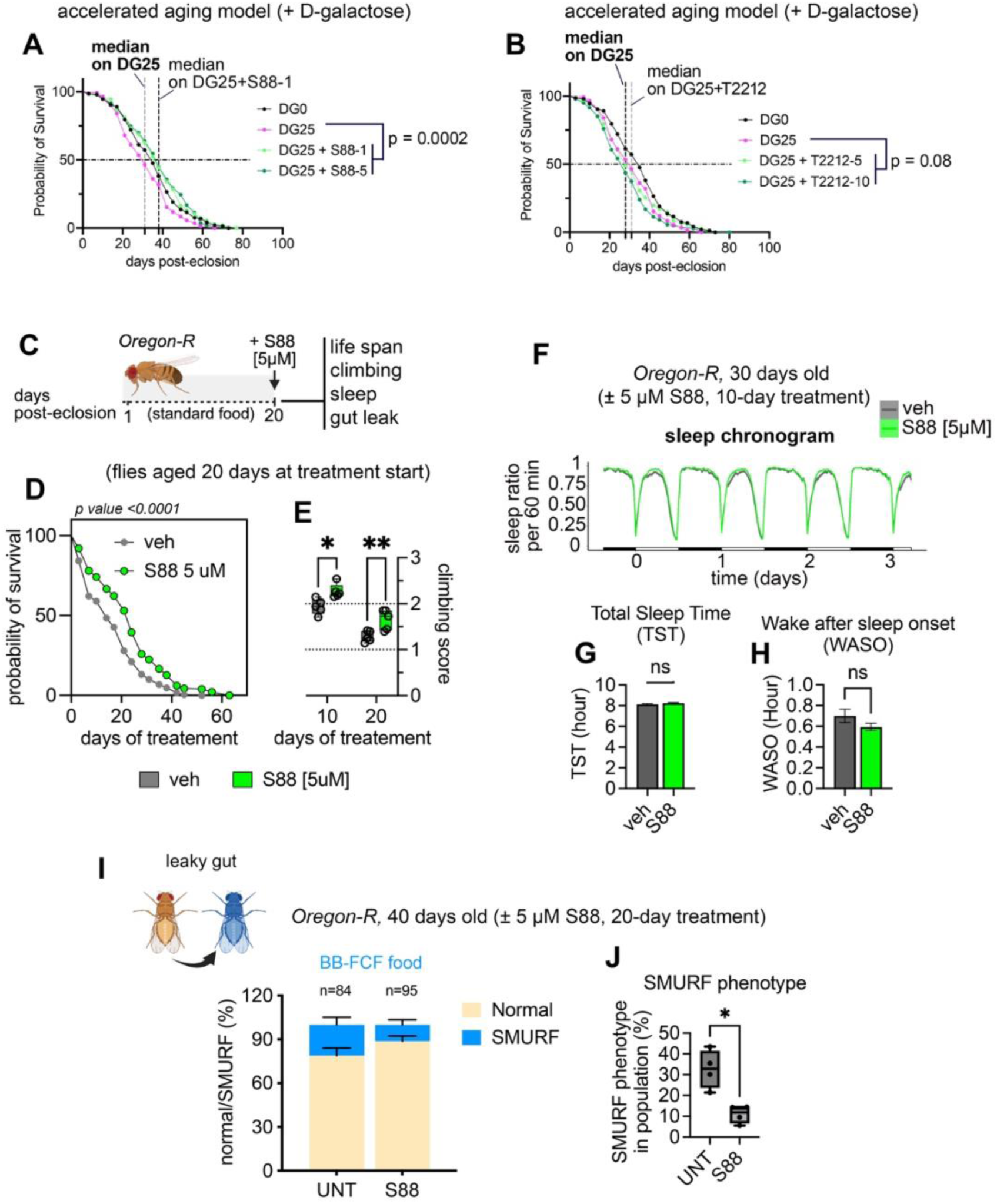
S88 improves aging-associated phenotypes in male *Drosophila*. (A, B) Survival of male flies subjected to a D-galactose–induced accelerated aging paradigm (DG25) and treated with S88 (1 or 5 µM; A), T2212 (1 or 5 µM; B), or left untreated. Survival of control flies maintained on standard food (DG0) is shown for reference. The median survival of the DG25 vehicle group is indicated (grey dotted line). n = 100 flies per condition. Statistical significance was determined by log-rank (Mantel–Cox) test. The dose-dependent effect of D-galactose on lifespan is shown in Fig. S13. (C) Experimental design for late-life intervention during physiological aging. Male flies were aged on standard food until day 20 post-eclosion and then transferred to food containing S88 (5 µM) or vehicle. Lifespan, climbing performance, locomotor activity/sleep parameters, and intestinal barrier function were assessed. (D) Survival following initiation of S88 treatment at day 20. n = 100 flies per condition. Statistical significance was determined by log-rank (Mantel–Cox) test. (E) Negative geotaxis performance after late-life S88 intervention. Climbing was assessed at days 30 and 40 (10 or 20 days of treatment, respectively). Statistical analysis was performed using one-way ANOVA with Šidák’s multiple-comparisons test at each age. (F) Representative sleep/activity chronogram of 30-day-old male flies recorded using the Drosophila Activity Monitoring System (DAMS) under a 12 h:12 h light–dark cycle over 3 days on S88 (5 µM) or vehicle food. (G, H) Sleep parameters quantified from the same 3-day DAMS recording, including total sleep time (TST) (G) and wake after sleep onset (WASO) (H). Comparisons were performed using unpaired t-tests. (I, J) Intestinal barrier integrity assessed by the Smurf assay in 40-day-old males following initiation of S88 treatment at day 20 (20 days of treatment). (I) Proportion of Smurf versus non-Smurf flies in vehicle and S88-treated groups. (J) Percentage of Smurf flies per biological replicate (n = 4 replicates, 25 flies per replicate), shown as box-and-whisker plots. Statistical analysis was performed using unpaired t-test.

To determine whether S88 also benefits physiological aging in the absence of exogenous pro-aging stressors, we next tested S88 in wild-type *Oregon-R* male flies using a late-life intervention paradigm. Flies were maintained on standard food until day 20 post-eclosion and then transferred to food containing S88 (5 µM) or vehicle (Fig. 7C). Initiation of S88 treatment at day 20 resulted in a significant extension of lifespan compared with vehicle-treated controls (Fig. 7D). Moreover, S88-treated flies displayed improved negative geotaxis performance at days 30 and 40 (corresponding to 10 and 20 days of treatment, respectively) (Fig. 7E).

Sleep becomes progressively disrupted with age in *Drosophila*, with reduced sleep consolidation and increased nocturnal arousals, as we have previously shown ^41^ mirroring age-associated sleep disruption observed across species, including humans ^42^. We therefore monitored sleep/activity profiles using the Drosophila Activity Monitoring System (DAMS) over three consecutive days under a 12 h:12 h light–dark cycle in 30-day-old flies that had received S88 treatment for 10 days (Figure 7F). Quantification of total sleep time (TST) and wake after sleep onset (WASO) revealed no significant change in TST following S88 treatment, while WASO showed a modest, non-significant trend toward reduction relative to vehicle-treated flies (Figure 7G, H).

Because age-dependent loss of intestinal barrier integrity (“gut leak”) is another established hallmark of physiological aging in *Drosophila* and is associated with systemic decline and increased mortality ^43^, we next assessed gut permeability using the Smurf assay ^44^ after 20 days of treatment (day 40). Under these conditions, S88-treated flies exhibited a reduced frequency of Smurf phenotypes relative to vehicle controls (Fig. 8I and J), consistent with improved maintenance of intestinal barrier function during aging.

**Figure 8.**
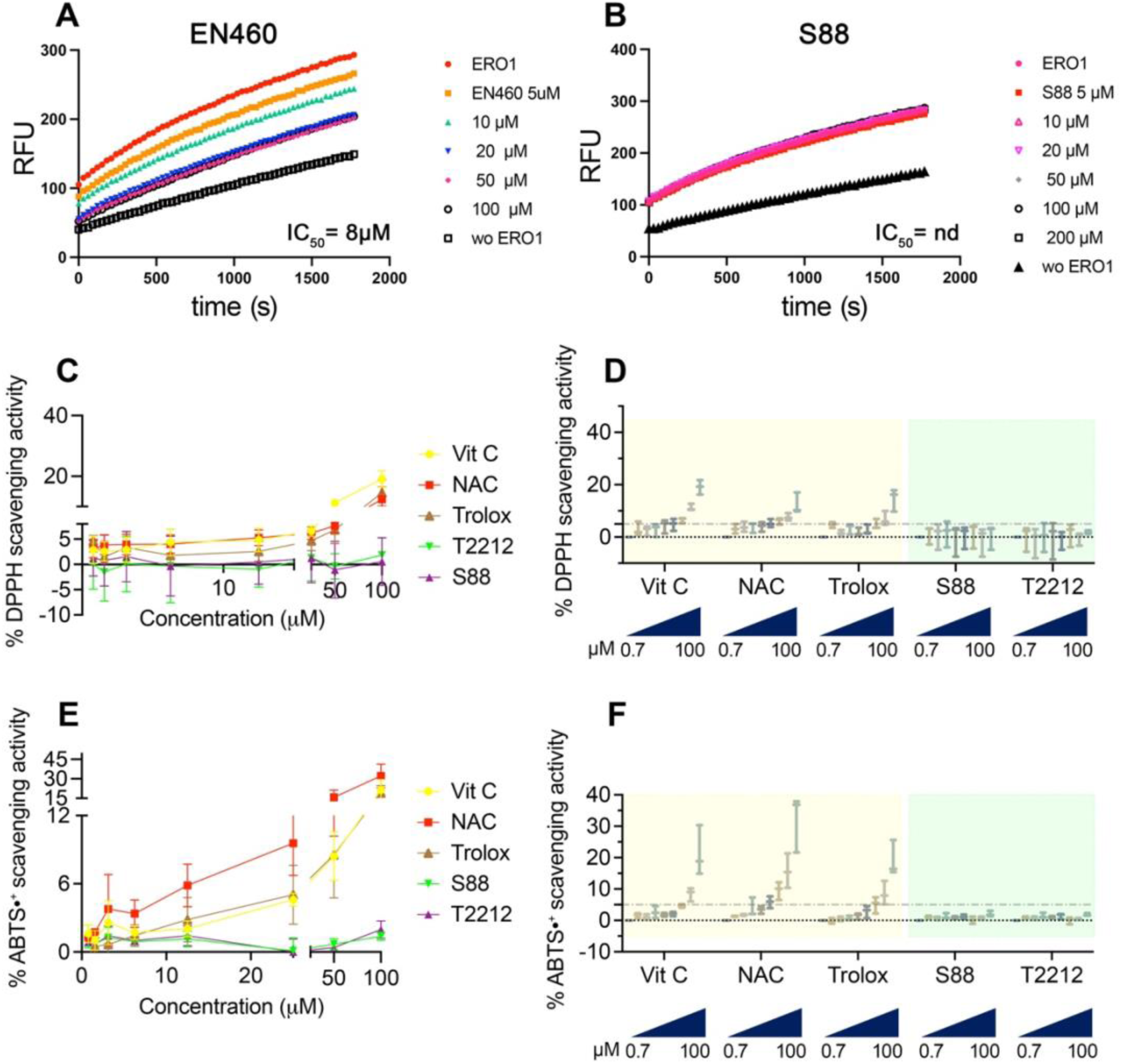
*In* vitro assays do not detect inhibition of ERO1A–PDIA1 activity or radical scavenging activity by S88. **(A–B)** Time course of Amplex UltraRed (AUR) fluorescence (RFU) in reactions containing recombinant ERO1A and PDIA1 in the presence of increasing concentrations of EN460 (A) or S88 (B). The control reaction containing ERO1α and PDIA1 alone is shown for comparison. An IC₅₀ (in µM) value was determined for EN460, whereas no measurable inhibition was detected for S88 under the conditions tested (nd, not detected). (C–D) DPPH radical scavenging activity of increasing concentrations (0.78125–100 μM) of S88 or T2212 compared with ascorbic acid (vitamin C), N-acetylcysteine (NAC), and Trolox as positive controls. (E–F) ABTS•⁺ radical scavenging activity of the same compounds under equivalent conditions. For clarity, data in (D) and (F) are presented as box-and-whisker plots (min to max). Data represent three independent experiments.

Together, these findings indicate that S88 confers benefits in both an accelerated aging paradigm and a late-life intervention model of physiological aging, supporting its potential to mitigate age-associated decline *in vivo*.

### 2.6 Biochemical assays do not support direct inhibition of ERO1A, PDI, or radical scavenging by S88

To further clarify the molecular basis of S88 activity, we sought to determine whether the compound directly inhibits the ER oxidase ERO1A, which initially guided compound prioritization during the structure-based screening pipeline. Molecular docking predicted a potential interaction of S88 within the ERO1A catalytic pocket; however, such predictions do not necessarily translate into functional inhibition in biochemical assays. To evaluate this possibility, we measured ERO1A activity using a coupled ERO1A–PDI oxidase assay. Under the conditions tested, S88 did not inhibit ERO1A activity (Fig. 8A and B). These results suggest that the protective phenotypes observed in cellular and *in vivo* models are unlikely to arise from direct inhibition of ERO1A or PDI.

Because cytoprotective effects can sometimes arise from general antioxidant properties, we additionally assessed radical scavenging activity and found no detectable direct scavenging by S88 (Fig. 8C-F).

Importantly, these findings do not exclude the possibility that S88 modulates ER-stress responses through indirect mechanisms or through interactions with other components of ER redox or proteostasis pathways that are not captured in simplified *in vitro* assays. Indeed, S88 consistently reduced ER-stress markers and promoted neuronal resilience in cellular models and improved neuromuscular and longevity phenotypes in *Drosophila* models of proteotoxic stress and aging.

Together, these results suggest that S88 acts as a modulator of ER-stress-associated pathways in physiological contexts, although its precise molecular target remains to be defined.

## 3. Discussion

Protein folding in the ER is tightly coupled to redox chemistry and oxygen radical production. This coupling supports efficient disulfide bond formation but also creates vulnerability: under proteostatic stress, oxidative folding can become a source of redox imbalance that exacerbates ER dysfunction. Such maladaptive UPR states, characterized by perturbed redox homeostasis and proteotoxicity, are increasingly recognized as convergent contributors to neurodegenerative disease and age-associated functional decline ^45-47^. From a therapeutic perspective, attenuation of ER stress therefore represents not only a means of alleviating downstream toxicity but also a strategy to interrupt self-reinforcing stress circuits that amplify cellular vulnerability.

Within this framework, the ER oxidase ERO1A has attracted particular attention because it sits at a mechanistic intersection between oxidative folding capacity and ER oxidative burden. By coupling disulfide bond formation to the production of hydrogen peroxide, ERO1A can shift from a homeostatic component of the folding machinery to a potential amplifier of oxidative stress. Consistent with this view, both genetic and pharmacological attenuation of ERO1A have been reported to reduce ER-stress outputs and improve proteostasis-linked phenotypes in multiple disease contexts ^48,49^. These observations provided a biologically grounded rationale to include ERO1A in our *in silico* prioritization framework. Accordingly, molecular docking was used as a hypothesis-generating step to evaluate potential compound-protein interactions, whereas compound advancement was determined primarily by phenotypic efficacy under ER-stress conditions.

A central conceptual outcome of this study is that effective ER-stress mitigation is best captured through convergent functional readouts rather than individual assays. Compounds can improve cell survival under stress without meaningfully reducing ER-stress signaling, while partial suppression of UPR markers does not necessarily translate into functional rescue. To address this, we implemented a prioritization strategy requiring both improved neuronal viability and reduction of ER-stress marker induction. This dual criterion enriched for candidates more likely to engage stress-relevant biology in a mechanistically interpretable manner. Within this framework, S88 consistently outperformed other candidates across neuronal assays and *in vivo* paradigms, suggesting that it modulates a process sufficiently central to influence both molecular stress markers and organismal function. Of note, T2212 (also known as geniposide) has previously been reported to possess neuroprotective properties and to modulate ER stress ^50,51^.

The divergence between S88 and other candidates is also informative. Several compounds displayed variable activity across experimental systems, which is common during early discovery and may reflect differences in cellular access, metabolic stability, or context-dependent engagement of stress pathways. More broadly, this variability underscores the biological complexity of ER-stress phenotypes, which are shaped by tissue-specific redox environments, proteostatic demand and compensatory signaling networks. The fact that S88 retains activity across human neurons, ALS-model flies and aging paradigms suggests that it influences a stress-relevant process with broad physiological leverage rather than a narrowly context-dependent endpoint.

In the UBQLN2^P497H^ *Drosophila* ALS model, S88 produced a sustained rescue of locomotor function and neuromuscular junction integrity, reaching levels comparable to healthy reference controls in the assays performed. Durable functional improvement is notable in chronic proteotoxic settings, where progressive tissue dysfunction and compensatory signaling often limit long-term efficacy. The accompanying reduction of ER-stress markers and selective transcriptional changes at later disease stages suggests that S88 may reshape the stress landscape during disease progression rather than producing a purely acute effect. Interestingly, ATF4 levels remained elevated despite reduced eIF2α phosphorylation and unchanged mTORC1 activity, raising the possibility that S88 promotes a non-canonical mode of ATF4 regulation. Because ATF4 participates in both adaptive and maladaptive stress programs, defining which downstream outputs remain active under S88 treatment may help clarify how ER-stress attenuation can coexist with preserved cellular resilience.

S88 also extended lifespan and improved healthspan-associated phenotypes in aging paradigms, including benefits in an accelerated D-galactose model and when administered later in life during physiological aging. These findings broaden the potential relevance of S88 beyond disease-specific contexts and support the view that ER-stress-associated redox imbalance may represent a modifiable contributor to organismal aging.

Importantly, S88 displayed minimal radical-scavenging activity in DPPH and ABTS assays, arguing against a simple antioxidant mechanism. Instead, these observations are more consistent with modulation of stress signaling or proteostasis-linked redox regulation, an important distinction given the frequent disconnect between *in vitro* antioxidant capacity and robust physiological benefit *in vivo*.

Biochemical analyses further indicated that S88 did not inhibit the activity of recombinant ERO1A in a typical coupled ERO1–PDI enzymatic assay. Thus, our data do not support direct enzymatic inhibition of ERO1A by S88 *in vitro*, and the proximate molecular target(s) responsible for its activity remain to be identified. However, the absence of inhibition in a purified biochemical assay does not exclude the possibility that S88 modulates ERO1A function in a cellular context. ER redox enzymes operate within highly compartmentalized and partner-dependent reaction networks, in which interactions with protein partners, local redox conditions, and subcellular organization may influence enzyme activity and regulatory outcomes compared with simplified in vitro assays. Moreover, phenotypic parallels with previously reported ERO1A attenuation and EN460-associated effects in the UBQLN2 ALS model ^11^ are consistent with the broader interpretation that S88 intersects with ER redox and ER-stress pathways along an ERO1-linked axis, even if the interaction is indirect or mediated through upstream regulators.

From a chemical standpoint, S88 contains a pyrazolopyridine scaffold that is increasingly explored in medicinal chemistry and compatible with diverse target classes ^52-54^. Related pyrazolone-based chemotypes have also gained attention in recent years, including FDA-approved drugs such as edaravone, which acts as a free-radical scavenger and is used in the treatment of amyotrophic lateral sclerosis (ALS). ^55^. Notably, pyrazolone-derived inhibitors targeting ERO1 have recently been reported in disease contexts characterized by dysregulated ER redox homeostasis, including ERO1-driven triple-negative breast cancer and SEPN1-related myopathy ^56^. While this feature makes S88 an attractive starting point for chemical optimization, it also highlights the importance of systematic target-deconvolution approaches, including proteome-wide engagement profiling and genetic interaction mapping, to distinguish primary binding partners from downstream signaling consequences. Such studies will be important to clarify the molecular basis of S88 activity, define biomarkers of target engagement and guide further optimization of this chemical series.

## 4. Conclusion

Using a multi-modal prioritization pipeline integrating structure-informed screening with functional validation across cellular and organismal models, we identified S88 as a natural-product lead that mitigates ER-stress-associated phenotypes. S88 improves neuromuscular function in UBQLN2^ALS^ *Drosophila*, extends lifespan in an accelerated aging model, and confers benefits during physiological aging, highlighting its capacity to promote resilience across ER-stress-linked contexts.

Together, these findings support two conclusions. First, attenuation of ER-stress signaling represents a tractable strategy to enhance neuronal and organismal resilience in disease and aging. Second, S88 emerges as a promising lead that improves ER-stress-associated molecular and functional phenotypes across models despite its molecular target(s) remaining unresolved. Defining how S88 interfaces with ER redox and stress-signaling networks, and whether it modulates ERO1A directly or indirectly, will be important for advancing this chemistry toward therapeutic development.

## 5. Materials and Methods

### 5.1 Phytochemical compound virtual screening

The three-dimensional (3D) structure of ERO1A was obtained from the Protein Data Bank (PDB) under accession ID 3AHQ, while the structure of ERO1B was predicted using SWISS-MODEL ^57^ using ERO1A as the template. The 3D structures of other FAD-containing enzymes, MAO-A, MAO-B, and LSD1, were retrieved from the PDB under accession IDs of 2Z5X, 2V5Z, and 6W4K, respectively. For all structures, non-protein molecules such as water, sodium, and magnesium ions were removed from the PDB files to retain only the protein molecules. Phytochemical compounds were obtained from the COCONUT database, consisting of 401,824 unique compounds. Initial filtering was based on drug-likeness properties, including a fraction of sp3 carbon atoms (Fsp3) > 0.4, lipophilicity (xlogp) < 5, and molecular weight (MW) < 500 Daltons. 3D structures (PDB format) of the passed compounds were generated from their SMILES format (smi) using Gypsum v1.2.0 ^58^. Python scripts, prepare_ligand4.py and prepare_receptor4.py from AutoDockTools ^59^, were used to convert PDB file of both ligands and protein targets into pdbqt format. A grid box was then set to cover the FAD-binding site on each protein target. Molecular docking simulations were conducted using AutoDock Vina ^60^ with an exhaustiveness setting of 8 to predict binding energy between ligands and protein targets. The lowest binding energy for each ligand was used to identify the most favorable docking pose. To evaluate the binding affinity of FAD-containing enzymes and their known inhibitors, the inhibitors from their original PDB files were removed and re-docked to their relevant protein targets using the same procedure. The binding affinity of these known inhibitors was used as a cutoff, and phytochemical compounds showing a binding affinity at least 1 kcal/mol lower than the inhibitors were considered to have favorable binding ability to the target. All scripts and their corresponding data are available on the executable platform at: https://github.com/asangphukieo/Phytochem_screening.

### 5.2 Compound handling and preparation

All chemicals were initially prepared as concentrated stock solutions in 100% DMSO and subsequently diluted to working concentrations, except for D-galactose (D-GAL). Retinoic acid (Sigma-Aldrich, CAS No. R2625) and tunicamycin (Sigma-Aldrich, CAS No. 11089-65-9) were prepared as 100 mM and 5 mg/mL stock solutions, respectively. The phytochemicals CNP0392367 (CAS No. STL522543), CNP0168092 (CAS No. STL551047), CNP0392367 (CAS No. STK179676), and CNP0162092 (CAS No. STL546888) were obtained from Vitas-M Laboratory (Hong Kong), and geniposide (CAS No. T2212) was purchased from TargetMol. All phytochemicals were prepared as 20 mM stock solutions in DMSO. Additional details on phytochemicals are provided in Supplementary Tables S1–S4. D-GAL was directly dissolved in the *Drosophila* culture medium at final concentrations of 25 mg/mL and 30 mg/mL. The compounds utilized in this study, along with their CAS registry numbers, are listed in Supplementary Table S7 unless otherwise specified.

### 5.3 Measurement of radical scavenging activity

The antioxidant capacity of selected phytochemicals was evaluated using two complementary assays that measure radical scavenging activity via different chemical mechanisms and solvent environments. The 1,1-diphenyl-2-picrylhydrazyl (DPPH) assay assesses the ability of compounds to neutralize lipophilic free radicals in methanolic solution, while the 2,2′-azino-bis(3-ethylbenzothiazoline-6-sulfonic acid) radical cation (ABTS) assay evaluates both hydrophilic and lipophilic antioxidant activity by measuring the reduction of a stable radical cation in aqueous conditions. For the DPPH assay, a 0.2 mM DPPH solution was freshly prepared in methanol. Test reactions were performed in 96-well plates according to the previously established protocol ^61^ by mixing 180 μl of the DPPH solution with 20 μl of each phytochemical dissolved in methanol at the indicated concentrations. Methanol alone (20 μl) was used as a negative control. Reactions were incubated at room temperature in the dark for 30 minutes, and absorbance was measured at 517 nm. Trolox, n-acetylcysteine and ascorbic acid served as positive controls. The percentage of DPPH radical scavenging was calculated after correcting for the intrinsic absorbance of the test compounds:

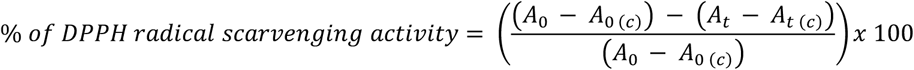

The absorbance of the control (A_0_) was corrected by subtracting the absorbance of methanol only (A_0 (c)_), and the absorbance of either tested phytochemicals or standards (A_t_) was corrected by subtracting the absorbance of methanol mixed with the tested compounds (A_t (c)_). Each compound was tested in triplicate across independent experiments (n ≥ 3).

For the ABTS assay, ABTS radical cations (ABTS•⁺) were generated by mixing equal volumes of 7 mM ABTS and 2.45 mM potassium persulfate, followed by incubation in the dark at room temperature for 12–16 hours, as described in ^62^. The resulting ABTS•⁺ solution was diluted 1:30 in Milli-Q water to achieve an absorbance of 0.70 ± 0.10 at 734 nm. In 96-well plates, 180 μl of diluted ABTS•⁺ solution was mixed with 20 μl of each test compound or control (dissolved in MQ water). After 6 minutes of incubation in the dark at room temperature, absorbance was measured at 734 nm. Trolox, n-acetylcysteine and ascorbic acid were used as positive controls. The percentage of scavenging inhibition was calculated using the following formula:

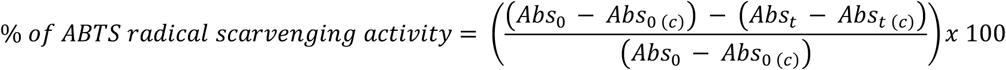

The absorbance of the control (A_0_) was corrected by subtracting the absorbance of methanol only (A_0 (c)_), and the absorbance of either tested phytochemicals or standards (A_t_) was corrected by subtracting the absorbance of methanol mixed with the tested compounds (A_t (c)_). All measurements were performed in triplicate and repeated in at least three independent experiments.

### 5.4 Human skin dermis-derived fibroblasts isolation

A skin biopsy was obtained from a healthy donor following a protocol approved by the Ethical Committee of the Faculty of Medicine, Chiang Mai University, under study code PED-2566-0380. A punch biopsy was collected and immediately placed in a 50 mL conical tube containing 10 mL of Dulbecco’s Modified Eagle Medium (DMEM; Gibco, Thermo Fisher Scientific, Waltham, MA, USA) supplemented with 1% penicillin-streptomycin (pen-strep; HyClone, Cytiva, Marlborough, MA, USA) to prevent contamination. Under sterile conditions, the biopsy was transferred into a digestion medium composed of DMEM, 20% fetal bovine serum (FBS; Sigma-Aldrich, St. Louis, MO, USA), collagenase type I (Gibco, Thermo Fisher Scientific, Cat# 17100-017), and 1% pen-strep. The tissue was incubated overnight at 37°C to facilitate enzymatic digestion. The following day, the sample was thoroughly mixed to promote epidermal separation and dermal disintegration. The mixture was then centrifuged to obtain a cell pellet, which was subsequently resuspended in fibroblast culture medium consisting of DMEM supplemented with 20% FBS and 1% pen-strep. The suspension was plated into a T75 culture flask and incubated at 37°C with 5% CO₂ for 6–7 days to allow fibroblast attachment. After 7 days, fibroblast-like cells were observed. At this stage, the culture medium was replaced with fibroblast culture medium containing 10% FBS to promote cell proliferation. Once the culture reached 80% confluency, cells were passaged at a 1:3 ratio using 0.25% trypsin-EDTA (HyClone, Cytiva, Marlborough, MA, USA). Human-derived fibroblasts (HDF) were expanded up to passage 3 and cryopreserved in liquid nitrogen for long-term storage.

### 5.5 Cell culture

Hepatocellular carcinoma (HepG2, ATCC: HB-8065) and neuroblastoma (SH-SY5Y, ATCC: CRL-2266) cell lines were obtained from the American Type Culture Collection (ATCC, Manassas, VA, USA). The HepG2 cells were cultured in T25 flasks using Dulbecco’s Modified Eagle Medium (DMEM) (Gibco, Grand Island, USA) supplemented with 10% fetal bovine serum (FBS) and 1% penicillin-streptomycin (both from Corning, USA). The SH-SY5Y cells were cultivated in T25 flasks using 1:1 mixture of Eagle’s Minimum Essential Medium (EMEM) (ATCC, Manassas, VA, USA) and F-12 medium (Gibco, Grand Island, USA), supplemented with 10% fetal bovine serum (Corning, Woodland, CA, USA) and 1% penicillin-streptomycin (Corning, Manassas, VA, USA). Cultures were maintained at 37°C with 5% CO₂, and all experiments were performed using cells at passages 6–12. All experiments involving cell culture were approved by the Institutional Biosafety Committee (ICB) of the Faculty of Medicine, Chiang Mai University (Thailand), under protocol number CMUIBC0268022.

### 5.6 Neuronal differentiation of SH-SY5Y cells

To induce neuronal differentiation, SH-SY5Y cells were seeded in 6-well plates at 1 × 10⁵ cells/well and cultured in EMEM/F-12 medium with 10% FBS for 24 hours. On day 0 (D0), differentiation was initiated by replacing the medium with differentiation medium containing 10 µM retinoic acid (RA) in EMEM/F-12 supplemented with 10% FBS. The RA-containing medium was refreshed every two days until day 5 (D5). Neuronal differentiation was confirmed by phase-contrast microscopy and Western blot analysis of neuronal marker expression.

### 5.7 Cell viability and cytotoxicity assay

Cell viability and cytotoxicity were assessed using the MTT assay (3-(4,5-dimethylthiazolyl-2)-2,5-diphenyltetrazolium bromide). The assay was used to evaluate (i) cell viability under tunicamycin (Tm)-induced ER stress, with or without phytochemical treatment, and (ii) the cytotoxic effects of the phytochemicals alone. For ER stress experiments, SH-SY5Y cells (either undifferentiated or differentiated with retinoic acid (RA) for 3 days) were used. Undifferentiated cells were seeded in 96-well plates at 2 × 10⁴ cells/well and cultured for 24h before treatment with Tm at 0.1, 1, or 10 µM for 24 or 48 hours. Differentiated cells were seeded at 3.3 × 10³ cells/well, maintained in RA until day 3, and treated with the same Tm concentrations and durations. To evaluate the cytotoxicity of the selected phytochemicals (S43, S47, S76, S88, and T2212), human dermal fibroblasts (HDF), HepG2, and SH-SY5Y cells were seeded in 96-well plates at 2.5 × 10³, 5 × 10³, and 2 × 10⁴ cells/well, respectively. After 24h of culture in EMEM/F12 medium, cells were treated with increasing concentrations (1, 2.5, 5, 10, and 20 µM) of each compound for 24 or 48h. To assess the potential protective effects of the phytochemicals against Tm-induced ER stress, SH-SY5Y cells (undifferentiated or 3-day RA-differentiated) were treated with 10 µM or 20 µM of each compound following 24 hours of Tm exposure. Vehicle-treated cells (0.1% DMSO) served as controls. For the MTT assay, culture media were replaced with 0.5 mg/mL MTT solution (BioChemica, Barcelona, Spain), and cells were incubated for 4h at 37 °C. The MTT-containing medium was then discarded, and 100 µL of DMSO (Labscan, Bangkok, Thailand) was added to dissolve the resulting formazan crystals. Absorbance was measured at 570 nm using a microplate reader (SpectraMax ABS Plus, MOLECULAR DEVICES). Unless otherwise specified, each condition included six technical replicates across two biological replicates. Results were expressed as a percentage of viable cells relative to controls. For untreated controls, viability was normalized to cells not exposed to DMSO. For Tm- and phytochemical-treated samples, viability was normalized to DMSO vehicle controls at either 0.05% or 0.1%, as appropriate.

### 5.8 *Drosophila* husbandry and stocks

*Drosophila melanogaster* stocks were maintained on a standard cornmeal-yeast-glucose medium at 22°C with 60% humidity under a 12:12 light/dark cycle unless otherwise stated. The Gal4/UAS binary system ^63^ was used for targeted, tissue-specific knockdown or overexpression. Pan-neuronal knockdown or overexpression was achieved using the nSyb-GAL4 or elav-GAL4 drivers at 25°C. The glass multiple reporter (GMR)-GAL4 promoter was used for targeting transgene’s expression to developing eyes at 28°C. The following transgenic fly lines were obtained from the Bloomington Drosophila Stock Center (BDSC): nSyb-GAL4 (#68222), elav-GAL4 (#8760), *UAS-dsRNA ERO1L* (#64674), *UAS-ERO1L* (#42698), *UAS-EGFP* (#5431), *UAS-AUG-DsRed* (#6282), *UAS-dsRNA green fluorescent protein* (*GFP IR*, #9330) and *UAS-HA-UBQLN2^P497H^* (hereafter referred to as *UBQLN2-ALS*) (#83363), UAS-LacZ (#1776). Wild-type *Oregon-R* flies were used as reference lineages (Bloomington #2376). To minimize genetic background effects, all experimental flies were backcrossed six times with the *w* strain before use.

### 5.9 *Drosophila* life span

The viability assay was performed as described in ^64^, with slight modifications. Virgin female flies carrying *nSyb-GAL4* (n = 10) were crossed with male flies carrying *UAS-ERO1L* (n = 10) on either phytochemical-supplemented food or standard medium containing 0.1% DMSO (vehicle control), with three replicates per condition. Phytochemicals were administered at concentrations of 1, 5, or 10 µM. Newly eclosed adult male flies overexpressing *ERO1L* were separated into low-density vials (20 flies per vial) containing either standard medium-vehicle or phytochemical-supplemented food. These flies were maintained under the same temperature and humidity conditions as described earlier. Flies were transferred to fresh food vials every three days, and mortality was recorded. All lifespan conditions were established in parallel within the same experimental batch under identical environmental conditions. A shared vehicle-treated control cohort was included as the reference population for comparisons across phytochemical treatments. Each phytochemical condition comprised 100 flies, distributed across 10 independent vials (10 flies per vial) to minimize vial-specific effects, whereas the shared control cohort comprised 389 flies to provide a robust baseline survival estimate. The survival rate was determined as the percentage of surviving flies each day until all flies had died. Comparisons were made between phytochemical-treated groups and the shared vehicle-treated control cohort. Survival curves were generated and analyzed using the Log-rank (Mantel-Cox) test for statistical validation.

### 5.10 *Drosophila* locomotive assays

Virgin female flies carrying *elav-GAL4* (n = 10) were crossed with males carrying *UAS-UBQLN2-ALS* (n = 10) on either standard medium or phytochemical-supplemented food, with three replicates per condition. S88 was provided at 1 or 5 µM, while T2212 was administered at 5 or 10 µM. Third instar larvae were selected for crawling assays, conducted as described in ^64^. Larval movement was tracked using ImageJ v1.53K with the wrMTrck plugin, and the distance traveled in 30 seconds was recorded. At least 55 larvae per group per condition were studied. Adult locomotion was assessed using a negative geotaxis assay as described in ^65^. Male flies from the same parental cross as described above were sorted and transferred to low-density vials (20 flies per vial) containing either standard medium or phytochemical-supplemented food. S88 was administered at 5 µM, and T2212 was administered at 10 µM. Flies aged 5, 10, 15, 20, 25, and 30 days post-eclosion were gently transferred, without anesthesia, into transparent conical tubes. The flies were acclimated for 10 minutes and then tapped to the bottom every 30 seconds before climbing (10 flies per vial, age-matched). Each trial was recorded and repeated five times. Climbing performance was scored based on the height reached within 5 seconds: 0 (<2.0 cm), 1 (2.0–3.9 cm), 2 (4.0–5.9 cm), 3 (6.0–7.9 cm), 4 (8.0–9.9 cm), and 5 (>10 cm). The climbing index was calculated as the weighted sum of the scores divided by the total number of flies per group. Scores were compared to age-matched controls, with 100 flies analyzed per genotype or treatment unless stated otherwise. Unless otherwise specified, a total of 100 flies per condition were studied.

### 5.11 Assessment of sleep parameters by *Drosophila* activity monitoring

Sleep parameters were assessed using Drosophila Activity Monitors (DAM; Trikinetics) as previously described ^41^, with minor modifications. Male flies were aged on standard food until day 30 post-eclosion and then transferred to monitoring tubes containing 5% glucose/1.5% agar food with or without S88 (5 µM) under a 12 h:12 h light–dark (LD) cycle. Sleep analysis was performed using 1-min bins and processed with Rtivity 1.2 software. A total of 96 flies were analyzed across three independent biological replicates. Sleep was defined as ≥5 min of uninterrupted inactivity. Total sleep time (TST) represents cumulative sleep during the dark phase (summed across the 3-day recording), and wake after sleep onset (WASO) denotes time spent awake during the dark phase following the first sleep bout (summed across the 3 - day recording).

### 5.12 Intestinal barrier integrity

Intestinal barrier integrity was assessed using the Smurf assay. Male wild-type *Oregon-R* flies were maintained on standard food until 20 days post-eclosion and then transferred to food containing S88 (5 µM) or vehicle for an additional 20 days. At 40 days of age, flies were transferred to Smurf dye food consisting of standard fly food supplemented with 0.1% (w/v) Brilliant Blue FCF (FH0202-A-GM025, Chemipan) and 5% sucrose, and allowed to feed for 18 h at 25 °C. Flies were then scored visually as “Smurf” (systemic blue coloration outside the digestive tract, indicating barrier leakage) or “non-Smurf” (blue coloration restricted to the gut). For each condition, 4 replicates were analyzed, with 25 flies per replicate (total n = 100 flies per condition). The percentage of Smurf flies per replicate was calculated and used for statistical analysis.

### 5.13 Neuromuscular junctions’ visualization

Third instar larvae expressing *UBQLN2^ALS^* in all neurons (elav-GAL4) were treated with 5 µM S88, 10 µM T2212, or left untreated (vehicle). The larvae were then dissected in hemolymph-like saline (HL3 saline), followed by fixation in 4% paraformaldehyde/PBS for 30 minutes, according to established methods ^66^. Blocking was carried out with a PBS solution containing 2% bovine serum albumin (BSA) and 0.1% Triton X-100. For detection, a fluorescein isothiocyanate (FITC)-conjugated goat anti-horseradish peroxidase antibody (1:1000, Jackson ImmunoResearch Laboratories, INC.) was used, and anti-Disc large (Dlg) mouse monoclonal antibody (4F3 anti-discs large, DSHB) was diluted to 1:300. The samples were mounted and examined using a confocal laser-scanning microscope (Olympus Fluoview FV3000). Quantification of glutamatergic motor neuron type I big (Ib) length and bouton counts in muscle 4 of abdominal segment 4 (A4) was performed. Image analysis was done with ImageJ (v. 1.53k). A total of 41, 45, and 40 NMJs from vehicle, S88-treated, and T2212-treated larvae, respectively, were analyzed.

### 5.14 RNA isolation and qPCR

Total RNA was isolated from undifferentiated SH-SY5Y cells using the RNeasy® Mini Kit (QIAGEN) following the manufacturer’s protocol. Three biological replicates were performed. RNA purity and concentration were assessed using a NanoDrop One/OneC Spectrophotometer (Thermo Fisher Scientific). Next, 1 µg of total RNA was reverse transcribed into cDNA with the SensiFAST cDNA Synthesis Kit (Bioline). Quantitative reverse transcription PCR (RT-qPCR) was carried out in triplicate for each RNA sample using the SensiFAST SYBR® Lo-ROX Kit on a CFX Opus Real-Time PCR System (Bio-Rad), with specific primers (Supplementary Table S5). Quantitative normalization for each sample was performed by using *ACTIN* as an internal control.

### 5.15 Proteins isolation and western blotting

Crude extracts were prepared from either third instar foraging larvae (L3) or adult *Drosophila* heads or RA-differentiated SH-SY5Y cells using RIPA Lysis and Extraction Buffer (Thermo Fisher), following the manufacturer’s instructions. Briefly, 50 L3 larvae or 40 *Drosophila* heads were homogenized in 400 µl or 100 µl of RIPA buffer supplemented with a protease inhibitor (Thermo Scientific™ Halt™ Protease Inhibitor Cocktail), respectively. SH-SY5Y cells were collected from 6-well plates, washed with PBS, and the cell pellet was resuspended in 60 µl of complete RIPA buffer. The homogenates were incubated at 4°C with gentle rotation for 20 minutes, then centrifuged at 14,000×g at 4°C for 20 minutes. The supernatant (cleared lysate) was collected, and protein concentration was determined using the BCA assay kit (Millipore #71285). A total of 30 µg of lysate was loaded onto a 12% polyacrylamide gel and transferred to an Immobilon-P PVDF membrane (Merck) using the Transblot-Turbo transfer system (Bio-Rad). The membranes were blocked with 5% skim milk in TBS-T (Tris-buffered saline with 0.1% Tween® 20) at room temperature and incubated overnight with primary antibodies (listed in Supplementary Table S6). After washing the membranes four times with TBS-T, they were incubated with HRP-conjugated secondary antibodies (1:3000) for 1 hour (Goat Anti-Rabbit or Anti-Mouse IgG (H + L)-HRP Conjugate, Bio-Rad #1706515 and #1706516, respectively). Primary and secondary antibodies were diluted in 5% skim milk in TBS-T. Membranes were developed using WesternSure Chemiluminescent Reagents (LI-COR) and imaged with the C-DiGit Blot Scanner (LI-COR). Protein levels were normalized to housekeeping proteins such as cytoplasmic Actin and nuclear Lamin C. For pEIF2α quantification, samples were normalized to total eIF2α protein levels. Band intensities were quantified using ImageJ software (version 1.53k) across multiple biological replicates.

### 5.16 Expression and purification of ERO1A recombinant protein

A plasmid encoding GST-SMT3-ERO1α was transformed into Rosetta (DE3) bacteria (Novagen). When the bacterial culture reached an optical density (OD600) between 0.6 and 0.8, expression was induced with 0.5 mM isopropyl-β-D-thiogalactoside (IPTG) at 16 °C for 24 hours. Bacterial cells were harvested by centrifugation and the pellet was resuspended in lysis buffer (200 mM NaCl, 50 mM Tris-HCl, pH 8.0, 0.2% Triton X-100) supplemented with 1 mM DTT, EDTA-free protease inhibitors, and DNase I, and the suspension was sonicated. After centrifugation for 30 minutes at 16,100 × g to remove insoluble material, the supernatant was collected and applied to a GST-Trap Glutathione Sepharose 4B column (GE17-0756-01, Merck) overnight at 4 °C. After three washes with high-salt buffer, Ulp protease (Z03691, GenScript) was added to the column and incubated for two hours at 4 °C. ERO1α was then collected, and its concentration was determined spectrophotometrically using a NanoDrop at 280 nm.

### 5.17 Expression and purification of PDIA1 recombinant protein

Recombinant His-tagged PDIA1 protein was expressed in Rosetta (DE3) bacteria (Novagen). When the bacterial culture reached an optical density (OD600) between 0.6 and 0.8, expression was induced with 0.5 mM isopropyl-β-D-thiogalactoside (IPTG) at 16 °C for 24 hours. Bacterial cells were harvested by centrifugation, and the pellet was resuspended in lysis buffer (200 mM NaCl, 50 mM phosphate buffer, pH 8.0, 0.2% Triton X-100) supplemented with EDTA-free protease inhibitors and DNase I, followed by sonication.After centrifugation for 30 minutes at 16,100 × g to remove insoluble material, the supernatant was collected and applied to a Ni²⁺ affinity column (Ni Sepharose™ 6 Fast Flow, Merck) overnight at 4 °C. After initial washes with 40 mM imidazole, PDIA1 was eluted with 250 mM imidazole and collected. Protein concentration was determined spectrophotometrically at 280 nm.

### 5.18 AUR-based kinetic assay to test ERO1A activity

The kinetic assay for ERO1α activity is based on a reaction involving recombinant ERO1α and PDIA1 proteins, horseradish peroxidase (Worthington), and 5 μM Amplex UltraRed (AUR; Invitrogen), as described in ^23^. PDI was previously reduced with DTT and desalted using a PD-10 column. The reaction was monitored kinetically at 535 nm excitation and 590 nm emission using a TECAN Infinite M Nano+ fluorescence plate reader. The effect of EN460 and S88 on ERO1α activity was evaluated by pre-incubating the compounds with ERO1α and measuring the inhibition of the linear reaction rate, followed by calculation of the corresponding IC₅₀ values using Prism 10 (GraphPad).

### 5.19 Data analysis

Statistical analyses were performed in GraphPad Prism 9 (GraphPad Software). Unless otherwise indicated, data are shown as mean ± SD, with individual data points displayed where feasible. For two-group comparisons, we used unpaired two-tailed Student’s t tests, applying Welch’s correction when variances were unequal. For experiments with more than two groups, we used one-way ANOVA with the multiple-comparisons procedure specified in the figure legend (Dunnett’s, Šidák’s, or Tukey’s). When data did not meet parametric assumptions, we applied nonparametric tests (Mann–Whitney for two groups; Kruskal–Wallis with Dunn’s post hoc test for multiple groups). Survival curves were compared using the log-rank (Mantel–Cox) test. Exact sample sizes (n), numbers of independent biological replicates, and the statistical tests used are reported in the corresponding figure legends. Statistical significance was defined as p < 0.05.

## Acknowledgments

This study was supported by Fundamental Fund, Chiang Mai University under the grants FF66/061, FF67/044 and FF68/207569 and supported by the Faculty of Medicine, Chiang Mai University grant no FACMED 139/2567. This study was partially supported by Genomics Thailand, Health Systems Research Institute (HSRI) under grants 66-124 and 68-060 and the by the Mid-Career Research Grant (N42A670768) of the National Research Council of Thailand. We thank Mr. Papon Hitmool, Mr. Sakorn Phongchankhiao, and Mr. Tawan Munwanna (Department of Pharmacology, Faculty of Medicine, Chiang Mai University), as well as Ms. Thunpitcha Meesawat and Ms. Ratchanaree Suepsaipaeng, (CMUTEAM, Faculty of Medicine, Chiang Mai University), for their technical and administrative assistance. We thank Dr. Yoshida Hideki (Kyoto Institute of Technology) and the Bloomington Stock Center for providing fly strains, the Science and Educational Company Limited (SCIED) for training in confocal scanning electron microscopy. Finally, we sincerely appreciate Dr. Natrujee Wiwattanadittakul, M.D. (Faculty of Medicine, Chiang Mai University), for collecting skin specimens for fibroblast isolation and extend our gratitude to the anonymous donor for the participation in this study.

## Author Contributions

Salinee Jantrapirom, Apiwat Sangphukieo and Luca Lo Piccolo designed the research. Salinee Jantrapirom, Apiwat Sangphukieo, Natsinee U-on and Luca Lo Piccolo carried out the experiments and performed data analysis. Pattaporn Poonsawas, Wasinee Wongkumool, Chansunee Panto, Puttachat Poound, Ranchana Yeewa, Ester Zito and Alice Marrazza participated part of the experiments. Salinee Jantrapirom, Apiwat Sangphukieo and Luca Lo Piccolo wrote the manuscript. All authors have reviewed and approved the final version of this manuscript.

## References

1 Lindholm, D., Wootz, H. & Korhonen, L. ER stress and neurodegenerative diseases. Cell Death Differ 13, 385–392 (2006). 10.1038/sj.cdd.4401778

2 Sprenkle, N. T., Sims, S. G., Sanchez, C. L. & Meares, G. P. Endoplasmic reticulum stress and inflammation in the central nervous system. Mol Neurodegener 12, 42 (2017). 10.1186/s13024-017-0183-y

3 Xiang, C., Wang, Y., Zhang, H. & Han, F. The role of endoplasmic reticulum stress in neurodegenerative disease. Apoptosis 22, 1–26 (2017). 10.1007/s10495-016-1296-4

4 Chen, X., Shi, C., He, M., Xiong, S. & Xia, X. Endoplasmic reticulum stress: molecular mechanism and therapeutic targets. Signal Transduct Target Ther 8, 352 (2023). 10.1038/s41392-023-01570-w

5 Pollard, M. G., Travers, K. J. & Weissman, J. S. Ero1p: a novel and ubiquitous protein with an essential role in oxidative protein folding in the endoplasmic reticulum. Mol Cell 1, 171–182 (1998). 10.1016/s1097-2765(00)80018-0

6 Zhang, J. et al. Endoplasmic Reticulum stress-dependent expression of ERO1L promotes aerobic glycolysis in Pancreatic Cancer. Theranostics 10, 8400–8414 (2020). 10.7150/thno.45124

7 Bassot, A. et al. The endoplasmic reticulum kinase PERK interacts with the oxidoreductase ERO1 to metabolically adapt mitochondria. Cell Rep 42, 111899 (2023). 10.1016/j.celrep.2022.111899

8 Haynes, C. M., Titus, E. A. & Cooper, A. A. Degradation of misfolded proteins prevents ER-derived oxidative stress and cell death. Mol Cell 15, 767–776 (2004). 10.1016/j.molcel.2004.08.025

9 Marciniak, S. J. et al. CHOP induces death by promoting protein synthesis and oxidation in the stressed endoplasmic reticulum. Genes Dev 18, 3066–3077 (2004). 10.1101/gad.1250704

10 Curran, S. P. & Ruvkun, G. Lifespan regulation by evolutionarily conserved genes essential for viability. PLoS Genet 3, e56 (2007). 10.1371/journal.pgen.0030056

11 Yeewa, R. et al. ERO1A inhibition mitigates neuronal ER stress and ameliorates UBQLN2(ALS) phenotypes in Drosophila melanogaster. Prog Neurobiol 242, 102674 (2024). 10.1016/j.pneurobio.2024.102674

12 Pagani, M. et al. Endoplasmic reticulum oxidoreductin 1-lbeta (ERO1-Lbeta), a human gene induced in the course of the unfolded protein response. J Biol Chem 275, 23685–23692 (2000). 10.1074/jbc.M003061200

13 Zito, E., Chin, K. T., Blais, J., Harding, H. P. & Ron, D. ERO1-beta, a pancreas-specific disulfide oxidase, promotes insulin biogenesis and glucose homeostasis. J Cell Biol 188, 821–832 (2010). 10.1083/jcb.200911086

14 Blais, J. D. et al. A small molecule inhibitor of endoplasmic reticulum oxidation 1 (ERO1) with selectively reversible thiol reactivity. J Biol Chem 285, 20993–21003 (2010). 10.1074/jbc.M110.126599

15 Voronkova, M. A. et al. ERO1A levels are a prognostic indicator in EGFR mutated non small cell lung cancer. NPJ Precis Oncol 8, 250 (2024). 10.1038/s41698-024-00736-1

16 Salvagno, C., Mandula, J. K., Rodriguez, P. C. & Cubillos-Ruiz, J. R. Decoding endoplasmic reticulum stress signals in cancer cells and antitumor immunity. Trends Cancer 8, 930–943 (2022). 10.1016/j.trecan.2022.06.006

17 Marciniak, S. J., Chambers, J. E. & Ron, D. Pharmacological targeting of endoplasmic reticulum stress in disease. Nat Rev Drug Discov 21, 115–140 (2022). 10.1038/s41573-021-00320-3

18 Germani, S. et al. SEPN1-related myopathy depends on the oxidoreductase ERO1A and is druggable with the chemical chaperone TUDCA. Cell Rep Med 5, 101439 (2024). 10.1016/j.xcrm.2024.101439

19 Hayes, K. E. et al. Inhibition of the FAD containing ER oxidoreductin 1 (Ero1) protein by EN-460 as a strategy for treatment of multiple myeloma. Bioorg Med Chem 27, 1479–1488 (2019). 10.1016/j.bmc.2019.02.016

20 Tu, B. P. & Weissman, J. S. The FAD- and O(2)-dependent reaction cycle of Ero1-mediated oxidative protein folding in the endoplasmic reticulum. Mol Cell 10, 983–994 (2002). 10.1016/s1097-2765(02)00696-2

21 Huijbers, M. M., Montersino, S., Westphal, A. H., Tischler, D. & van Berkel, W. J. Flavin dependent monooxygenases. Arch Biochem Biophys 544, 2–17 (2014). 10.1016/j.abb.2013.12.005

22 Johnson, B. D. et al. Identification of Natural Product Sulfuretin Derivatives as Inhibitors for the Endoplasmic Reticulum Redox Protein ERO1alpha. ACS Bio Med Chem Au 2, 161–170 (2022). 10.1021/acsbiomedchemau.1c00062

23 Varone, E. et al. Small molecule-mediated inhibition of the oxidoreductase ERO1A restrains aggressive breast cancer by impairing VEGF and PD-L1 in the tumor microenvironment. Cell Death Dis 16, 105 (2025). 10.1038/s41419-025-07426-1

24 Atanasov, A. G., Zotchev, S. B., Dirsch, V. M., International Natural Product Sciences, T. & Supuran, C. T. Natural products in drug discovery: advances and opportunities. Nat Rev Drug Discov 20, 200–216 (2021). 10.1038/s41573-020-00114-z

25 Xie, S., Zhan, F., Zhu, J., Xu, S. & Xu, J. The latest advances with natural products in drug discovery and opportunities for the future: a 2025 update. Expert Opin Drug Discov 20, 827–843 (2025). 10.1080/17460441.2025.2507382

26 Ryoo, H. D. Drosophila as a model for unfolded protein response research. BMB Rep 48, 445–453 (2015). 10.5483/bmbrep.2015.48.8.099

27 Yan, C. et al. IRE1 promotes neurodegeneration through autophagy-dependent neuron death in the Drosophila model of Parkinson’s disease. Cell Death Dis 10, 800 (2019). 10.1038/s41419-019-2039-6

28 Katsube, H., Hinami, Y., Yamazoe, T. & Inoue, Y. H. Endoplasmic reticulum stress-induced cellular dysfunction and cell death in insulin-producing cells results in diabetes-like phenotypes in Drosophila. Biol Open 8 (2019). 10.1242/bio.046524

29 Deng, H. X. et al. Mutations in UBQLN2 cause dominant X-linked juvenile and adult-onset ALS and ALS/dementia. Nature 477, 211–215 (2011). 10.1038/nature10353

30 Wu, J. J. et al. ALS/FTD mutations in UBQLN2 impede autophagy by reducing autophagosome acidification through loss of function. Proc Natl Acad Sci U S A 117, 15230–15241 (2020). 10.1073/pnas.1917371117

31 Halloran, M. et al. Amyotrophic lateral sclerosis-linked UBQLN2 mutants inhibit endoplasmic reticulum to Golgi transport, leading to Golgi fragmentation and ER stress. Cell Mol Life Sci 77, 3859–3873 (2020). 10.1007/s00018-019-03394-w

32 Renaud, L., Picher-Martel, V., Codron, P. & Julien, J. P. Key role of UBQLN2 in pathogenesis of amyotrophic lateral sclerosis and frontotemporal dementia. Acta Neuropathol Commun 7, 103 (2019). 10.1186/s40478-019-0758-7

33 Torrence, M. E. et al. The mTORC1-mediated activation of ATF4 promotes protein and glutathione synthesis downstream of growth signals. Elife 10 (2021). 10.7554/eLife.63326

34 Ben-Sahra, I., Hoxhaj, G., Ricoult, S. J. H., Asara, J. M. & Manning, B. D. mTORC1 induces purine synthesis through control of the mitochondrial tetrahydrofolate cycle. Science 351, 728–733 (2016). 10.1126/science.aad0489

35 Hetz, C. & Dillin, A. Central role of the ER proteostasis network in healthy aging. Trends Cell Biol (2024). 10.1016/j.tcb.2024.10.003

36 Lopez-Otin, C., Blasco, M. A., Partridge, L., Serrano, M. & Kroemer, G. The hallmarks of aging. Cell 153, 1194–1217 (2013). 10.1016/j.cell.2013.05.039

37 Bhattarai, K. R., Riaz, T. A., Kim, H. R. & Chae, H. J. The aftermath of the interplay between the endoplasmic reticulum stress response and redox signaling. Exp Mol Med 53, 151–167 (2021). 10.1038/s12276-021-00560-8

38 Qiao, X. et al. ER reductive stress caused by Ero1alpha S-nitrosation accelerates senescence. Free Radic Biol Med 180, 165–178 (2022). 10.1016/j.freeradbiomed.2022.01.006

39 Azman, K. F. & Zakaria, R. D-Galactose-induced accelerated aging model: an overview. Biogerontology 20, 763–782 (2019). 10.1007/s10522-019-09837-y

40 Shwe, T., Pratchayasakul, W., Chattipakorn, N. & Chattipakorn, S. C. Role of D-galactose-induced brain aging and its potential used for therapeutic interventions. Exp Gerontol 101, 13–36 (2018). 10.1016/j.exger.2017.10.029

41 Yeewa, R. et al. The histone acylation reader ENL/AF9 regulates aging in Drosophila melanogaster. Neurobiol Aging 144, 153–162 (2024). 10.1016/j.neurobiolaging.2024.10.002

42 Pai, S. K. Mechanisms underlying fragmented sleep in aging. Aging Brain 3, 100077 (2023). 10.1016/j.nbas.2023.100077

43 Salazar, A. M., Aparicio, R., Clark, R. I., Rera, M. & Walker, D. W. Intestinal barrier dysfunction: an evolutionarily conserved hallmark of aging. Dis Model Mech 16 (2023). 10.1242/dmm.049969

44 Rera, M., Clark, R. I. & Walker, D. W. Intestinal barrier dysfunction links metabolic and inflammatory markers of aging to death in Drosophila. Proc Natl Acad Sci U S A 109, 21528–21533 (2012). 10.1073/pnas.1215849110

45 Ajoolabady, A., Lindholm, D., Ren, J. & Pratico, D. ER stress and UPR in Alzheimer’s disease: mechanisms, pathogenesis, treatments. Cell Death Dis 13, 706 (2022). 10.1038/s41419-022-05153-5

46 Meares, G. P. et al. PERK-dependent activation of JAK1 and STAT3 contributes to endoplasmic reticulum stress-induced inflammation. Mol Cell Biol 34, 3911–3925 (2014). 10.1128/MCB.00980-14

47 Bellani, S. et al. GRP78 clustering at the cell surface of neurons transduces the action of exogenous alpha-synuclein. Cell Death Differ 21, 1971–1983 (2014). 10.1038/cdd.2014.111

48 Shergalis, A. G., Hu, S., Bankhead, A., 3rd & Neamati, N. Role of the ERO1-PDI interaction in oxidative protein folding and disease. Pharmacol Ther 210, 107525 (2020). 10.1016/j.pharmthera.2020.107525

49 Jha, V. et al. ERO1-PDI Redox Signaling in Health and Disease. Antioxid Redox Signal 35, 1093–1115 (2021). 10.1089/ars.2021.0018

50 Yamazaki, M., Chiba, K. & Yoshikawa, C. Genipin suppresses A23187-induced cytotoxicity in neuro2a cells. Biol Pharm Bull 32, 1043–1046 (2009). 10.1248/bpb.32.1043

51 Ma, Z. G. et al. Protection against cardiac hypertrophy by geniposide involves the GLP-1 receptor / AMPKalpha signalling pathway. Br J Pharmacol 173, 1502–1516 (2016). 10.1111/bph.13449

52 Atukuri, D. Pyrazolopyridine: An efficient pharmacophore in recent drug design and development. Chem Biol Drug Des 100, 376–388 (2022). 10.1111/cbdd.14098

53 Waly, O. M., El-Sayed, S. M., Ghaly, M. A. & El-Subbagh, H. I. Multi-targeted anti-Alzheimer’s agents: Synthesis, biological evaluation, and molecular modeling study of some pyrazolopyridine hybrids. Eur J Med Chem 262, 115880 (2023). 10.1016/j.ejmech.2023.115880

54 Zhao, Z. et al. Pyrazolone structural motif in medicinal chemistry: Retrospect and prospect. Eur J Med Chem 186, 111893 (2020). 10.1016/j.ejmech.2019.111893

55 Lapchak, P. A. A critical assessment of edaravone acute ischemic stroke efficacy trials: is edaravone an effective neuroprotective therapy? Expert Opin Pharmacother 11, 1753–1763 (2010). 10.1517/14656566.2010.493558

56 Retini, M. et al. Pyrazolone-based ERO1 inhibitors in ERO1-driven triple-negative breast cancer and SEPN1-related myopathy: Structure-activity relationship and therapeutic potential. Pharmacol Res 222, 108037 (2025). 10.1016/j.phrs.2025.108037

57 Waterhouse, A. et al. SWISS-MODEL: homology modelling of protein structures and complexes. Nucleic Acids Res 46, W296–W303 (2018). 10.1093/nar/gky427

58 Ropp, P. J. et al. Gypsum-DL: an open-source program for preparing small-molecule libraries for structure-based virtual screening. J Cheminform 11, 34 (2019). 10.1186/s13321-019-0358-3

59 Morris, G. M. et al. AutoDock4 and AutoDockTools4: Automated docking with selective receptor flexibility. J Comput Chem 30, 2785–2791 (2009). 10.1002/jcc.21256

60 Trott, O. & Olson, A. J. AutoDock Vina: improving the speed and accuracy of docking with a new scoring function, efficient optimization, and multithreading. J Comput Chem 31, 455–461 (2010). 10.1002/jcc.21334

61 Kiriya, C., Yeewa, R., Khanaree, C. & Chewonarin, T. Purple rice extract inhibits testosterone-induced rat prostatic hyperplasia and growth of human prostate cancer cell line by reduction of androgen receptor activation. J Food Biochem 43, e12987 (2019). 10.1111/jfbc.12987

62 Re, R. et al. Antioxidant activity applying an improved ABTS radical cation decolorization assay. Free Radic Biol Med 26, 1231–1237 (1999). 10.1016/s0891-5849(98)00315-3

63 Brand, A. H. & Perrimon, N. Targeted gene expression as a means of altering cell fates and generating dominant phenotypes. Development 118, 401–415 (1993). 10.1242/dev.118.2.401

64 Lo Piccolo, L., et al. FAME4-associating YEATS2 knockdown impairs dopaminergic synaptic integrity and leads to seizure-like behaviours in Drosophila melanogaster. Prog Neurobiol 233, 102558 (2024). 10.1016/j.pneurobio.2023.102558

65 Jantrapirom, S., Lo Piccolo, L., Yoshida, H. & Yamaguchi, M. Depletion of Ubiquilin induces an augmentation in soluble ubiquitinated Drosophila TDP-43 to drive neurotoxicity in the fly. Biochim Biophys Acta Mol Basis Dis 1864, 3038–3049 (2018). 10.1016/j.bbadis.2018.06.017

66 Lo Piccolo, L. & Yamaguchi, M. RNAi of arcRNA hsromega affects sub-cellular localization of Drosophila FUS to drive neurodiseases. Exp Neurol 292, 125–134 (2017). 10.1016/j.expneurol.2017.03.011

